# Muscle length modulates recurrent inhibition and post-activation depression differently according to contraction type

**DOI:** 10.1101/2024.12.10.627478

**Authors:** Julian Colard, Julien Duclay, Yohan Betus, Thomas Cattagni, Marc Jubeau

**Affiliations:** Nantes Université, Movement - Interactions - Performance, MIP, UR 4334, F-44000 Nantes, France; Toulouse NeuroImaging Center, Université de Toulouse, Inserm, UPS, Toulouse, France

**Author notes:** CORRESPONDING AUTHOR Marc Jubeau Nantes University, Movement - Interactions - Performance, MIP, UR 4334, F-44000 Nantes, France 25 bis Boulevard Guy Mollet - BP 72206 44 322 Nantes cedex 3, France.

**Keywords:** eccentric, long muscle length, Ia afferent, H reflex, Renshaw cell, GABA

## Abstract

It is well documented that, in soleus, motoneuron output and the effectiveness of activated Ia afferents to discharge α-motoneurons both decrease during eccentric contractions. Evidence suggests that these regulations can be explained by (1) recurrent inhibition and (2) greater post-activation depression by primary afferent depolarization. However, the influence of muscle length on the regulation of the effectiveness of Ia afferents to discharge α-motoneurons observed during eccentric contractions remains unclear. We conducted a study on 16 healthy young individuals. We used simple and conditioned Hoffmann reflex with different conditioning techniques such as paired H reflex, D1 method and heteronymous Ia facilitation coupled with electromyography during eccentric, isometric and concentric contractions at long, intermediate and short soleus muscle lengths. Our results confirm that during eccentric contraction the effectiveness of Ia afferents to discharge _U_-motoneurons decreases only at intermediate and short muscle lengths but is similar between all contraction types at long muscle length. Findings are similar for recurrent inhibition. Post-activation depression is significantly more pronounced during eccentric contractions compared with isometric and concentric contractions at long muscle length. Our analysis also shows that recurrent inhibition and post-activation depression are greater at long muscle length compared with short muscle length, whatever the contraction type. These new findings demonstrate an important influence of muscle length on the activity of spinal regulatory mechanisms and the effectiveness of activated Ia afferents to discharge α-motoneurons during eccentric contractions.

## INTRODUCTION

Motoneuron output and muscle activation are generally less during eccentric contractions than during isometric and concentric contractions (Westing *et al*., 1991; Komi *et al*., 2000; Pasquet *et al*., 2000; Aagaard *et al*., 2000; Babault *et al*., 2003). Investigations into corticospinal and spinal pathways reveal a specific neural control during eccentric contractions (Duclay *et al*., 2011, 2014; Duchateau & Enoka, 2016), which is thought to come mainly from a significant inhibition of the regulatory mechanisms acting at the spinal level (Gruber *et al*., 2009; Howatson *et al*., 2011; Duclay *et al*., 2014).

At the spinal level, the effectiveness of activated Ia afferents to discharge α-motoneurons is reduced during submaximal or maximal eccentric contractions in the soleus (SOL) (Duclay & Martin, 2005). Among the possible inhibitory mechanisms, the contribution of the recurrent inhibition generated by Renshaw cell activity (Barrué-Belou *et al*., 2018, 2019; Papitsa *et al*., 2022) and post-activation depression by primary afferent depolarization (PAD, (Colard *et al*., 2023) have been demonstrated. Recurrent inhibition and post-activation depression by PAD are modulated by both supraspinal control (Haase & van der meulen, 1961; Grosprêtre *et al*., 2014) and peripheral pathways (Hultborn *et al*., 1979; Rudomin & Schmidt, 1999). During eccentric contractions, both of these mechanisms decrease the SOL H reflex but coexist without influencing each other (Papitsa *et al*., 2022).

It is important to note that these observations, i.e., greater recurrent inhibition and post-activation depression by PAD, were obtained from measurements on the SOL muscle in the neutral ankle position (0°), which corresponds to an intermediate muscle length. However, muscle length could influence these processes. Indeed, changes in muscle length that occur during dynamic actions, especially eccentric contractions, might also activate type I (dynamic and static sensitivities) and II (only static) mechanoreceptor afferent discharges projecting to the spinal cord (Matthews, 2011; Dimitriou, 2022). Furthermore, another study demonstrated that contractile elements of the medial gastrocnemius muscle, such as sarcomeres or fascicles, undergo specific extreme stretching during the final phases of eccentric contractions (Guilhem *et al*., 2016). Consequently, the mechanical stress occurring within the muscle-tendon unit during eccentric contractions exerts differential influences upon receptors that are sensitive to muscle length (e.g. muscle spindles) or tension (e.g. Golgi tendon organs), depending on the joint position. One original finding was that, at long muscle length, voluntary activation level is less during eccentric contractions than during isometric and concentric ones (Doguet *et al*., 2017*b*). Moreover, at long muscle lengths during eccentric contractions, corticospinal excitability is comparable to that observed during isometric and concentric contractions (Doguet *et al*., 2017*a*), whereas at the same length, an increase in the silent period was noted during eccentric contractions compared to isometric and concentric ones. Although caution is necessary in interpreting silent period measurements (Kidgell *et al*., 2017), it is more likely that the deficits in voluntary activation and the modulations in corticospinal excitability observed during eccentric contractions at long muscle lengths are attributable to spinal rather than intracortical changes. The spinal mechanisms involved in modulation during eccentric contractions are post-activation depression by PAD and recurrent inhibition. It is possible that these mechanisms are modulated differently at greater muscle lengths. Our previous results have shown that post-activation depression by PAD is more pronounced during passive muscle lengthening at long muscle lengths (Colard *et al*., 2024). These results are very interesting and allow us to speculate that there could be an increase in post-activation depression by PAD during eccentric contractions at long muscle lengths. Furthermore, the _U_-motoneuron discharge rate is lower at longer muscle lengths than at shorter ones (Pasquet *et al*., 2006) and this reduction is closely linked to Renshaw cell activity (McCrea *et al*., 1980). It is therefore possible that Renshaw cells provide a greater inhibition during eccentric contractions at long muscle lengths. The effects of the interaction between muscle length and contraction type on the H reflex, recurrent inhibition and PAD activities remain to be elucidated.

This study aimed to explore the influence of muscle length on the effectiveness of activated Ia afferents to discharge α-motoneurons, as well as the associated pre- and post-synaptic inhibitory mechanisms, including recurrent inhibition and post-activation depression by PAD, during eccentric, isometric and concentric contractions at short, intermediate and long muscle lengths in SOL. We hypothesized that the heightened activity of recurrent inhibition (Barrué-Belou *et al*., 2018, 2019) and post-activation depression by PAD (Colard *et al*., 2023) observed during eccentric contractions would be more pronounced when these contractions occurred at longer muscle lengths in SOL.

## METHODS

### Participants

To ensure data accuracy, G*Power software (version 3.1.9.6) was used to determine whether the size of our test sample was adequate to obtain representative results for the variables of interest. This evaluation was conducted after data collection.

The sample size for H reflex testing was estimated using an effect size (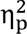 = 0.580) derived from the contraction type × muscle length interaction. With a desired statistical power (1 − β) of 0.95 and a significance level (α) of 0.05, a sample size of 8 participants was considered sufficient for a repeated measures ANOVA. The sample sizes for recurrent inhibition, post-activation depression by PAD, and heteronymous facilitation (HF) were estimated using effect sizes (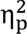 = 0.765, 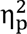 = 0.836, and ^2^= 0.923, respectively) calculated from the significant contraction type × muscle length interaction. With a power of 0.95 and an α of 0.05, the required sample sizes for these analyses were determined to be 8, 6, and 6 participants, respectively, for a repeated measures ANOVA.

A total of 16 participants (8 males and 8 females; age: 23 ± 2 years; height: 175 ± 4 cm; weight: 70 ± 13 kg) were included in this study. Among these individuals, 12 (6 males and 6 females; age: 21 ± 3 years; height: 172 ± 3 cm; weight: 67 ± 3 kg) voluntarily participated in a fourth session (see details below). Participants had no history of neurological injuries or diseases and provided written informed consent to participate. They were instructed to refrain from engaging in strenuous exercise for 48 hours prior to the testing sessions. Approval for the project was obtained from the Ethics Committee on Non-Interventional Research of Nantes University (n°12102021). All procedures conducted in this study met the requirements of the Declaration of Helsinki (last modified in 2013).

### Study design

All 16 participants took part in at least three separate experiments, with 12 of them agreeing to a fourth experimental session. Each experiment was separated by a minimum of two days. In each experiment, assessments of contraction type (eccentric, isometric and concentric contractions) and muscle length (short, intermediate and long muscle lengths) were randomly assigned to avoid potential biases stemming from order of testing procedures. In the first experiment (experiment A), assessments of soleus (SOL) fascicle length, muscle torque, H reflex and M wave recruitment curves in SOL were conducted. In the second experiment (experiment B), the same effects on recurrent inhibition were explored with paired H reflex techniques. The third experiment (experiment C) then investigated post-activation depression by PAD with the D1 method. Heteronymous Ia facilitation was assessed in an additional fourth experiment (experiment D). The order of sessions B and C was randomized among participants. All experimental data were collected from the participants’ right legs.

### Experimental set-up

#### Mechanical data

The participants were seated with the trunk inclined at 30°, the hip flexed at 85° (0° representing anatomical position) and the knee joint extended at 0° (representing full extension). Torque values were collected using an isokinetic dynamometer (Biodex 3, Shirley, NY, USA). A Biodex ergometer enabled instantaneous recording of muscle torque at constant angular velocity (20 deg.s^−1^). The output signal from the dynamometer was collected at 2 kHz using a commercial acquisition system (CED power 1401-3A, Cambridge Electronic Design; Cambridge, UK), displayed and then stored with Signal 7 software (Cambridge Electronic Design; Cambridge, UK). The foot of the right leg was strapped to the footplate, while the trunk and thighs were secured to the seat of the isokinetic dynamometer. The ankle angle was modulated according to the experimental conditions required. During isometric contraction, ankle angle was set at +15° (plantarflexion), 0° (neutral position) and -15° (dorsiflexion), for short, intermediate and long muscle lengths, respectively. During dynamic contractions, the ankle angle range of motion was set at 30°. For eccentric contraction, ankle angle moved from 30° to 0°, 15° to -15° and 0° to -30° for short, intermediate and long muscle lengths, respectively. For concentric contraction, the ankle angle was changed from -30° to 0°, -15° to 15° and 0° to 30°. During the four experiments, the dynamometer movement cycles lasted 5 s, including 1.5 s for performing the movement (plantarflexion or dorsiflexion), 1.5 s to return to the initial position, and a 1-s rest before the next cycle. Additionally, before each measurement, the subjects performed a submaximal isometric plantarflexion (for 1 s) to obtain similar thixotropic effects between different test conditions.

#### Electromyography

Electromyograms were recorded on SOL, medial gastrocnemius, tibialis anterior and vastus lateralis muscles using pairs of self-adhesive surface electrodes (Meditrace 100; Covidien, Mansfield, MA, USA) in bipolar configuration with a 30 mm inter-electrode distance. For SOL, electrodes were placed 2 cm below the muscle–tendon junction of the gastrocnemii. For medial gastrocnemius, electrodes were fixed lengthwise over the middle of the muscle belly. For the tibialis anterior, electrodes were placed on the muscle belly parallel to the longitudinal axis of the muscle, one-third of the distance between the head of the fibula and the tip of the medial malleolus. For the vastus lateralis, electrodes were placed at a position two-thirds of the way along the line from the anterior spina iliac superior to the lateral side of the patella. Before fixing the electrodes, the skin was shaved, gently abraded and then cleaned with alcohol. Signals were amplified (1000×) with a g.BSamp 0201a bio-amplifier (Guger Technologies, Shieldberg, Austria) and bandpass filtered (5–500 Hz). The signals were sampled at 5 kHz with a data acquisition system (CED power 1401-3A, Cambridge Electronic Design; Cambridge UK) and acquired with Signal software (version 7; CED).

#### Percutaneous electrical nerve stimulation

The electrophysiological responses (i.e. H reflex and M wave) of the SOL were evoked by percutaneous electrical stimulation of the posterior tibial nerve with a single rectangular pulse (1 ms) and high voltage (400 V), delivered by a stimulator (Digitimer, model DS8R Biphasic Constant Current Stimulator, Hertfordshire, UK). A self-adhesive electrode (1-cm diameter, Ag-AgCl) was used as a cathode and was placed in the popliteal fossa. The anode electrode (5 × 10 cm, Medicompex SA, Ecublens, Switzerland) was placed on the anterior surface of the knee, below the patella. Once the position was determined, the cathode electrode was firmly fixed to this site with a strap and tape, as recommended by Cattagni *et al*. (2018). To study post-activation depression by PAD modulations (D1 method), the fibular nerve was stimulated with a DS7R Current Stimulator, (Digitimer, Hertfordshire, UK) by placing the cathode electrode close to the head of the fibula and the anode electrode near the medial part of the tibial head. For heteronymous facilitation, the femoral nerve was stimulated with the cathode positioned over the nerve in the femoral triangle, and the anode placed over the greater trochanter. The location of stimulation electrode was identified to obtain an M wave in the vastus lateralis associated with an upward movement of the patella. The motor thresholds for stimulation of the fibular and femoral nerves were determined as the lowest intensity that evoked at least three M waves out of five stimulations.

### Experimental protocol

#### Experiment A

Experiment A was carried out in two stages. In the first part, three maximal voluntary isometric contractions at 0° were first performed to determine the maximal SOL EMG_RMS_ level. The maximum SOL EMG activity recorded during isometric MVCs was used to determine the target activation level (50% of maximal EMG_RMS_) provided to the participants during the study. Participants were asked to contract their plantarflexors so that they moved and maintained their SOL EMG_RMS_ biofeedback on the target corresponding to 50% of maximal EMG_RMS_. The biofeedback corresponded to the root mean square (RMS) value of the EMG signal provided by a digital computing channel. This channel instantaneously computed the RMS level of the amplified EMG signal with an integration time of 500 ms. This method made it possible to keep levels of muscle activation constant between all test conditions (Duclay *et al*., 2014; Colard *et al*., 2023).

Then, in the second stage torque values (at 50% of maximal SOL EMG_RMS_) were collected. Subsequently, the SOL fascicle length was measured using a Mach30 ultrasound scanner (Hologic-Supersonic Imagine, Aix-en-Provence, France) with B-mode image acquisition and linear probe (SL18-5, 7.5 MHz central frequency, Hologic-Supersonic Imagine, Aix en Provence, France). Participants sat on the seat of the isokinetic dynamometer with their trunk inclined 20° backwards from the vertical and knee fully extended. The lateral malleolus was aligned with the axis of rotation of the dynamometer. Images were taken from +15° (plantarflexion) to -15° (dorsiflexion) for eccentric submaximal contraction, with 0° being the anatomical position. Fascicle length changes were evaluated during sub-maximal eccentric contractions (50% of maximal EMG_RMS_) with a constant angular velocity of 20 deg.s^−1^. We identified the proximal and distal insertions of the SOL muscle, positioning the probe just distal to the myotendinous junction of the medial gastrocnemius muscle due to the clear visibility of SOL fascicles in this area. Despite the SOL muscle’s multipennate structure, a previous study on its three-dimensional architecture showed no significant differences in fascicle length across its four compartments, i.e., medial-anterior, lateral-anterior, medial-posterior and lateral-posterior, (Bolsterlee *et al*., 2018). Our chosen probe position therefore provided a fascicle length measurement representative of the entire SOL muscle. The probe, secured in a foam holder, was firmly attached to the leg with tape. To condition the muscle, we performed five ankle angle rotations ranging from 15° (plantarflexion) to -15° (dorsiflexion) back and forth (Nordez *et al*., 2006). B-mode ultrasound images were continuously recorded through three acquisitions during eccentric contractions at 50% of maximal EMG_RMS_, covering the full range of motion, to assess the architecture of the SOL muscle. The acquired ultrasound images were sampled at 50 Hz and processed using a customized MATLAB script (Fig. 1).

**Figure 1.**
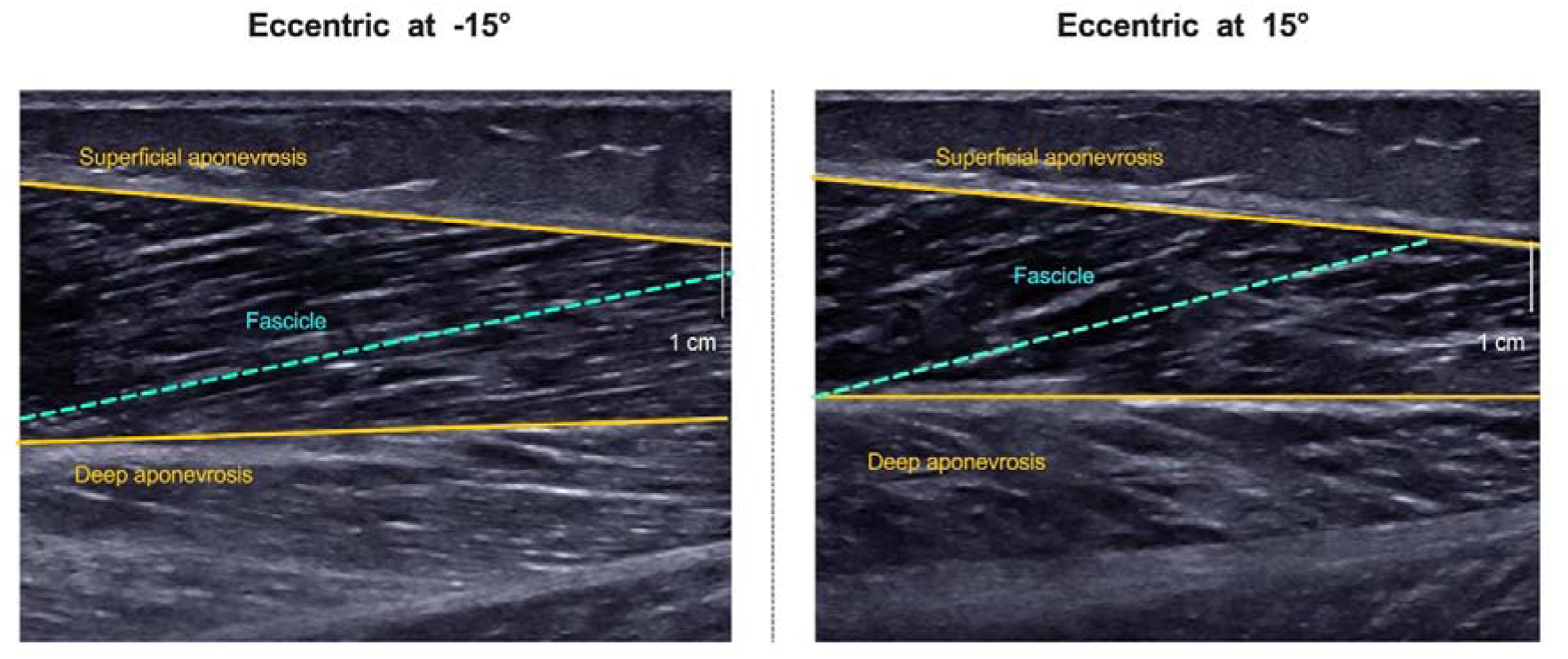
Examples of two ultrasound images in the sagittal plane during eccentric contraction at different ankle angles in soleus. This image was used to calculate soleus fascicle length (blue dashed line) from the visible insertion of the fibre between the deep and superficial aponeurosis (yellow lines).

During the second part of the experiment, maximal H reflex (H_max_) and maximal M wave (M_max_) were measured for all contraction types (eccentric, isometric and concentric) at each muscle length (short, intermediate, and long). SOL recruitment curves were determined during plantarflexion at 50% of maximal EMG_RMS_, resulting in a total of nine recruitment curves (see Fig. 2). The stimulation intensity was gradually increased with a 2-mA increment from the H reflex threshold to the intensity at which no further increase in the SOL M wave amplitudes was observed (i.e. a plateau). Finally, supramaximal stimulations at 150% of this latter stimulation intensity were delivered to ensure the recording of SOL M_max_. Five electrical nerve stimulations were delivered at each stimulation intensity. An interstimulus interval of 5 s was respected between each electrical nerve stimulation, as recommended in Hopkins *et al*. (2000) and Theodosiadou *et al*. (2023).

**Figure 2.**
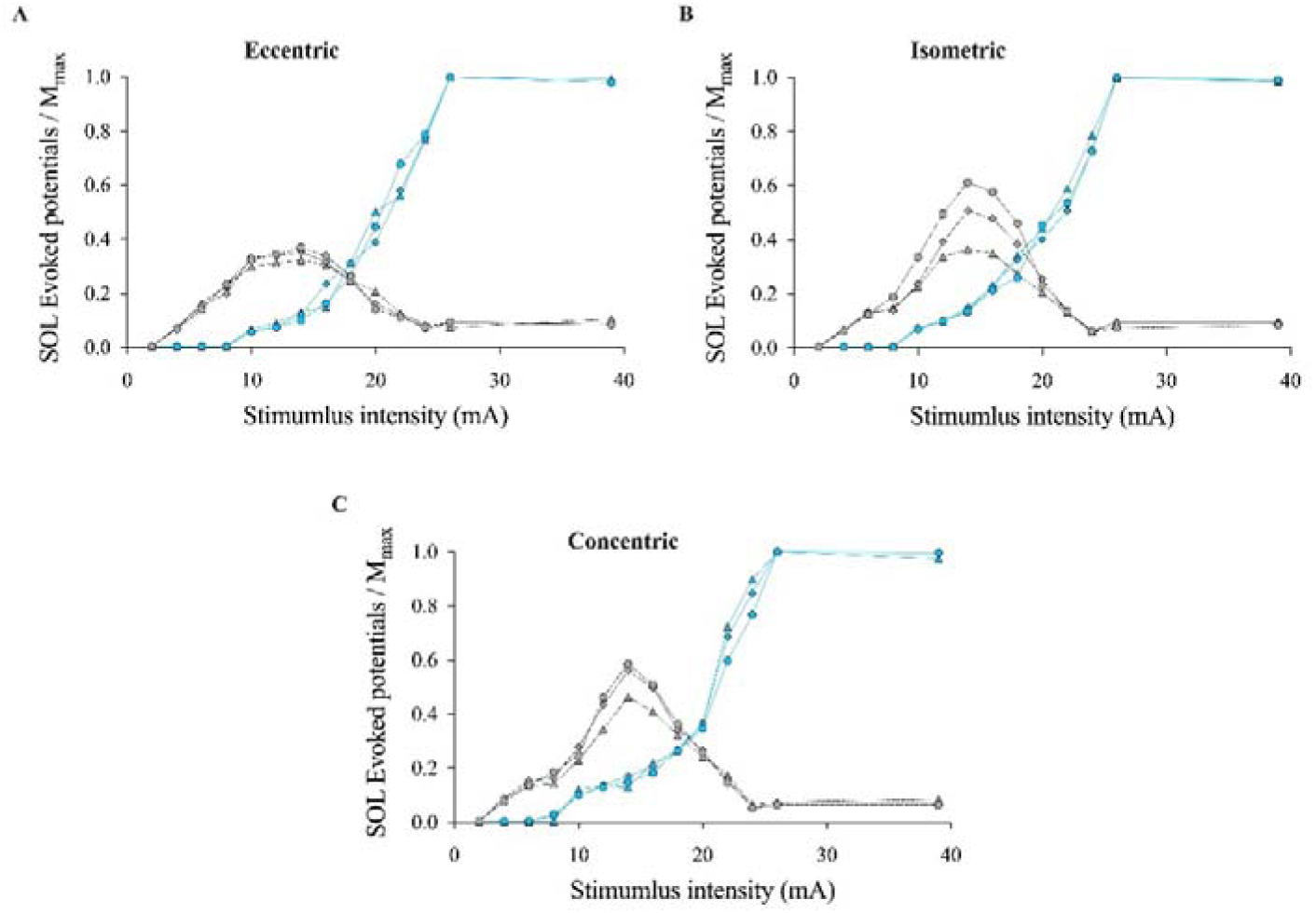
Recruitment curves of soleus H reflex and M wave according to muscle length and contraction type in one representative participant. Short (Δ Δ), intermediate (●, ●) and long (▴, ▴) recruitments curves of soleus H reflex (grey colour) and M wave (blue colour) during eccentric (A), isometric (B) and concentric (C) contraction for one representative participant.

Electrical tibial nerve stimulations were delivered at ankle angles of 15°, 0° and -15°, (the middle of each range of motion) for short, intermediate and long muscle lengths, respectively. Tibial nerve stimulations were delivered using the CED acquisition system. During eccentric and concentric contractions, stimulations were automatically delivered when the ankle angle passed +15° (short), 0° (intermediate) or -15° (long).

#### Experiment B

To assess recurrent inhibition, we employed the paired H reflex method developed by Pierrot-Deseilligny & Bussel (1975), in which two electrical stimuli are delivered over the posterior tibial nerve.

To assess the conditioning H reflex H_1_, four stimulations were delivered during each of four eccentric, concentric and isometric contractions at long, intermediate and short muscle lengths. The determination of H_1_ involved measuring the maximal H reflex without the presence of an M wave for each of the nine combinations of experimental conditions (Fig. 3). Prior to testing each condition, SOL H_1_ was assessed and fine-tuned to ensure that no M wave was present during the H_1_ assessment (Barrué-Belou *et al*., 2018). Renshaw cells, responsible for recurrent inhibition, are activated orthodromically by the H_1_ volley via recurrent collaterals, but they can also be activated antidromically by the direct activation of motor axons, which produce the M wave associated with H_1_. By optimising H_1_, we increased sensitivity to recurrent inhibition modulation while avoiding overestimation caused by the M wave. Four stimulations were delivered at supramaximal intensity generating M_max_ and V responses in the SOL muscle.

**Figure 3.**
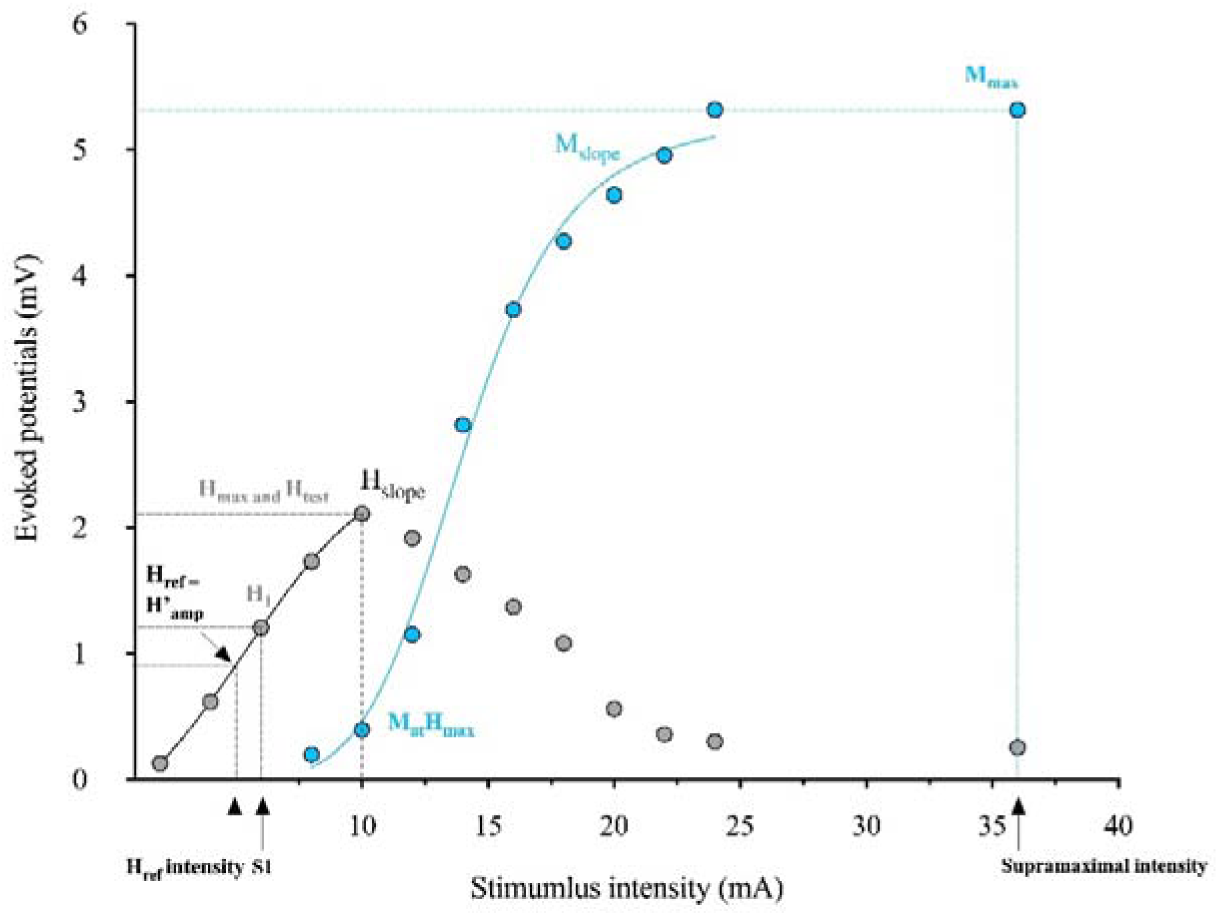
Representation of sigmoidal fittings on soleus recruitment curve of H reflex. 333 **(grey circles) and M wave (bleu circles), to determine optimal amplitudes and stimulation intensities for recurrent inhibition and post-activation by PAD assessments for one representative participant**. Hmax, maximal H reflex; Htest, test H reflex for post-activation by PAD assessment; H1, conditioning H reflex for recurrent inhibition; S1, stimulation intensity to evoked H1; Href, reference H reflex for recurrent inhibition; Href intensity, stimulation intensity to evoked Href; Mmax, maximal M wave; MatHmax, M wave corresponding at Hmax; supramaximal intensity, stimulation intensity to evoke Mmax. Hslope, H reflex slope development; Mslope M wave slope development.

To assess test H reflex (H’), eight paired H reflex (S_1_ + supramaximal intensity) stimulations were delivered with a 10-ms interval. The first stimulus (S1) was set at the intensity used to evoke the conditioning H reflex (H_1_), followed by a second stimulus at supramaximal intensity. The decrease observed between the conditioning H reflex (H_1_ evoked with S_1_ only) and the test H reflex (H’ evoked with S_1_ + supramaximal intensity) was attributed to recurrent inhibition (Hultborn *et al*., 1979).

Additional trials were recorded if the coefficient of variation of H_1_, M_max_ or H’ exceeded 5% across trials. To ensure accuracy and reliability of our results, we conducted the evaluation of each condition in a randomized order, with a 5-minute rest period between each.

#### Experiment C

To investigate post-activation depression by PAD activity, we used the D1 method, as described by Mizuno et al. (1971). This technique involves eliciting an electrical volley on the nerve that supplies the _a_-motoneuron pool of the antagonist muscle, thereby activating the Ia terminals and leading to a depression of the H reflex (Pierrot-Deseilligny & Burke, 2005). The extent of H reflex depression was previously believed to depend on the excitability of PAD interneurons, which could decrease the release of neurotransmitters at the Ia afferent terminals. SOL H_test_ was conditioned with a stimulus applied to the fibular nerve, which activates GABAergic interneurons responsible for primary afferent depolarization of the Ia afferents from the SOL muscle. To evoke SOL conditioned H reflex (H_D1_), a train of three 1-ms stimulations at 300 Hz was applied to the fibular nerve, with an intensity equivalent to 120% the stimulation intensity of the tibialis anterior motor threshold. The interval between the first shock of the train and the SOL H_test_ was 21 ms (Aymard *et al*., 2000; Lamy *et al*., 2009; Magalhães *et al*., 2015). Here, a total of 20 H_D1_ and 20 H_test_ were randomly evoked in all test conditions (eccentric, isometric and concentric contractions at short, intermediate and long muscle lengths).

#### Experiment D

To study heteronymous Ia facilitation, we used the method of Hultborn *et al*. (1987), which evaluates the monosynaptic facilitatory effect of femoral nerve stimulation on the amplitude of the SOL H reflex. Recent discoveries regarding the functioning of PAD indicate that afferent projections from the quadriceps to the SOL motoneuron are not directly inhibited by GABAergic interneurons but are consistently facilitated by them (employing the same mechanisms as PAD). However, heightened activity of these interneurons generates PAD-evoked spikes (Metz *et al*., 2023*a*) that can result in greater depletion of neurotransmitters, diminishing the facilitation reaching the SOL motoneuron and ultimately reducing H reflex facilitation. To minimize contamination from polysynaptic inputs, the femoral nerve stimulation needs to follow tibial nerve stimulation due to the shorter neural pathway of the heteronymous Ia afferent pathway compared with the homonymous pathway. By considering the mean duration of a synaptic event (0.5 to 1 ms) and systematically assessing facilitation onset with 1 ms steps, we aimed to minimize the influence of polysynaptic excitatory inputs (Baudry & Enoka, 2009; Johannsson *et al*., 2015; Souron *et al*., 2019). For each participant, the onset of H reflex facilitation in response to femoral nerve stimulation was determined by adjusting the delay between test (tibial nerve) and conditioning (femoral nerve) stimuli in 1-ms increments, ranging from 9 ms (test stimulus preceding conditioning stimulus by 9 ms) to 1 ms (test stimulus preceding conditioning stimulus by 1 ms). Conditioning stimulation intensity was set at 130% of the motor threshold, and an average of at least 5 trials was made for each delay. The criterion used to identify facilitation onset was an increase in conditioned H reflex (H_Fac_) amplitude of more than 10% over non-conditioned H reflexes (Johannsson *et al*., 2015). Totals of 20 H_Fac_ and 20 H_test_ were randomly evoked in all test conditions (eccentric, isometric and concentric contractions at short, intermediate and long muscle lengths).

To determine non-conditioned H reflexes, H_test_ was determined based on the smallest measurement of H_max_ amplitude among the nine experimental conditions (Fig. 3). This approach ensured that they were all subject to the same size of H_test_, allowing accurate comparisons to be made among them. Prior to testing, SOL H_test_ was adjusted to ensure that the same proportion of motor units contributing to the non-conditioned H reflexes were activated across all experimental conditions (Crone & Nielsen, 1989; Baudry & Duchateau, 2012). We conducted the different evaluations in a randomized order, with a 5-minute rest period between each.

Finally, two maximal voluntary eccentric, concentric and isometric dorsiflexions were performed at the ends of experiments A, B, C and D to evaluate the antagonist coactivation (3 seconds of duration, interspaced by 1-minute rest).

## Data analysis

### Evoked potentials

For each contraction type and muscle length, the mean peak-to-peak amplitude of SOL H_max_, M_max_, H_1_, H’, H_ref_, V, H_D1,_ H_Fac_ and H_test_ were calculated over all trials. All ratios of evoked potentials were calculated during eccentric, isometric and concentric contractions at short, intermediate and long muscle lengths.

To estimate the effectiveness of activated Ia afferents to discharge _a_-motoneurons we calculated H_max_/M_max_. This ratio is commonly used as an indicator for the maximal proportion of α-motoneurons recruited by the Ia afferent pathway. H_max_/M_max_ may be susceptible to perturbation by fluctuations in the M wave, primarily attributed to peripheral factors. In addition, the slope of the ascending limb of the recruitment curve reflects the rate of motoneuron recruitment as a function of the increase in Ia input to the motoneuron pool and is generally used to assess the reflex gain (Funase *et al*., 1994; Baudry *et al*., 2014; Penzer *et al*., 2015). This reflex gain is a complementary measure that specifically examines the gain and will therefore be more closely linked to transmission efficiency. In addition, by employing this strategy, the impact of peripheral factors on the results is minimized, ensuring accuracy and reliability of results (Hwang et al., 2002; Funase et al., 1994). Since sigmoidal analysis remains the most relevant method for analysing the H reflex development curve (Klimstra & Zehr, 2008), the sigmoidal slopes for H_slope_ and M_slope_ were determined by fitting the reflex development components to the ascendent part of the individual H and M recruitment curves (Fig. 3).

The M wave elicited concomitantly with the H_max_ (M_at_H_max_), a small fraction of the M_max_ (M_at_H_max_/M_max_), was measured and analysed to verify the stability of the stimulus intensity in each corresponding condition (Schieppati, 1987).

To check for experiment-induced changes in background motoneuron excitability, we selected the smallest H’ amplitude across all conditions. We then extrapolated the stimulus intensity required to obtain H’ amplitude without conditioning (H_ref_ intensity) through sigmoidal analysis on the corresponding recruitment curve. By plotting this stimulus intensity on all recruitment curves (using sigmoidal fitting), we extrapolated the corresponding H_ref_ amplitude for each condition (Fig. 3). If a greater variation occurred in H’ than in H_ref_, while the amplitude of the H_1_ remained constant, this indicated a change in recurrent inhibition. Conversely, if both reflexes varied in the same direction, no conclusions could be drawn regarding recurrent inhibition (Katz & Pierrot-Deseilligny, 1999).

To examine recurrent inhibition with the paired H reflex technique, we calculated H’/H_1_: A higher H’/H_1_ ratio is associated with a lower recurrent inhibition. H’/M_max_, H_1_/M_max_, V/M_max_ and H_ref_/M_max_ were calculated to ensure that the methodological criteria of paired H reflex had been respected.

To examine post-activation by PAD with the D1 method, we calculated H_D1_/H_test_. A higher H_D1_/H_test_ ratio is associated with a lower post-activation depression by PAD. H_test_/M_max_ was used to check that the amplitude of the non-conditioned H reflex was consistent between the different experimental conditions.

To examine heteronymous Ia facilitation with the heteronymous Ia facilitation method, we calculated H_Fac_/H_test_. A higher H_Fac_/H_test_ ratio is associated with a greater facilitation. H_test_/M_max_ was used to check that the amplitude of the non-conditioned H reflex was consistent between experimental conditions.

### EMG activity during voluntary contractions

A root mean square (RMS) of the SOL and medial gastrocnemius EMG signal over a 500-ms period prior to stimulation was normalized to the corresponding amplitude of M_max_ (EMG_RMS_/M_max_) during all conditions. The normalization procedure was used to ascertain whether SOL (agonist) and medial gastrocnemius (synergistic) EMG activities remained constant across all experimental conditions. Similarly, an RMS of the tibialis anterior EMG signal was used over the same time period preceding the stimulation to check that the coactivation was constant. To assess the level of coactivation in our different conditions, RMS activity of the tibialis anterior muscle was expressed relative to its activity during maximal isometric voluntary dorsiflexions for isometric contractions. For concentric trials, tibialis anterior RMS during concentric plantarflexions was normalized to maximal eccentric dorsiflexion. Similarly, for eccentric trials, tibialis anterior RMS during eccentric plantarflexions was normalized to maximal concentric dorsiflexion. The average level of coactivation was then calculated across all trials for each condition (Hagood *et al*., 1990).

### Muscle torque

Torque, produced by the plantarflexors during excentric submaximal contraction (50% of maximal EMG_RMS_), respectively, was noted at +15° (plantarflexion), 0° (neutral position) and -15° (dorsiflexion). The plantarflexor torque was computed as the mean value recorded over all trials for each ankle angle (Fig. 4).

**Figure 4.**
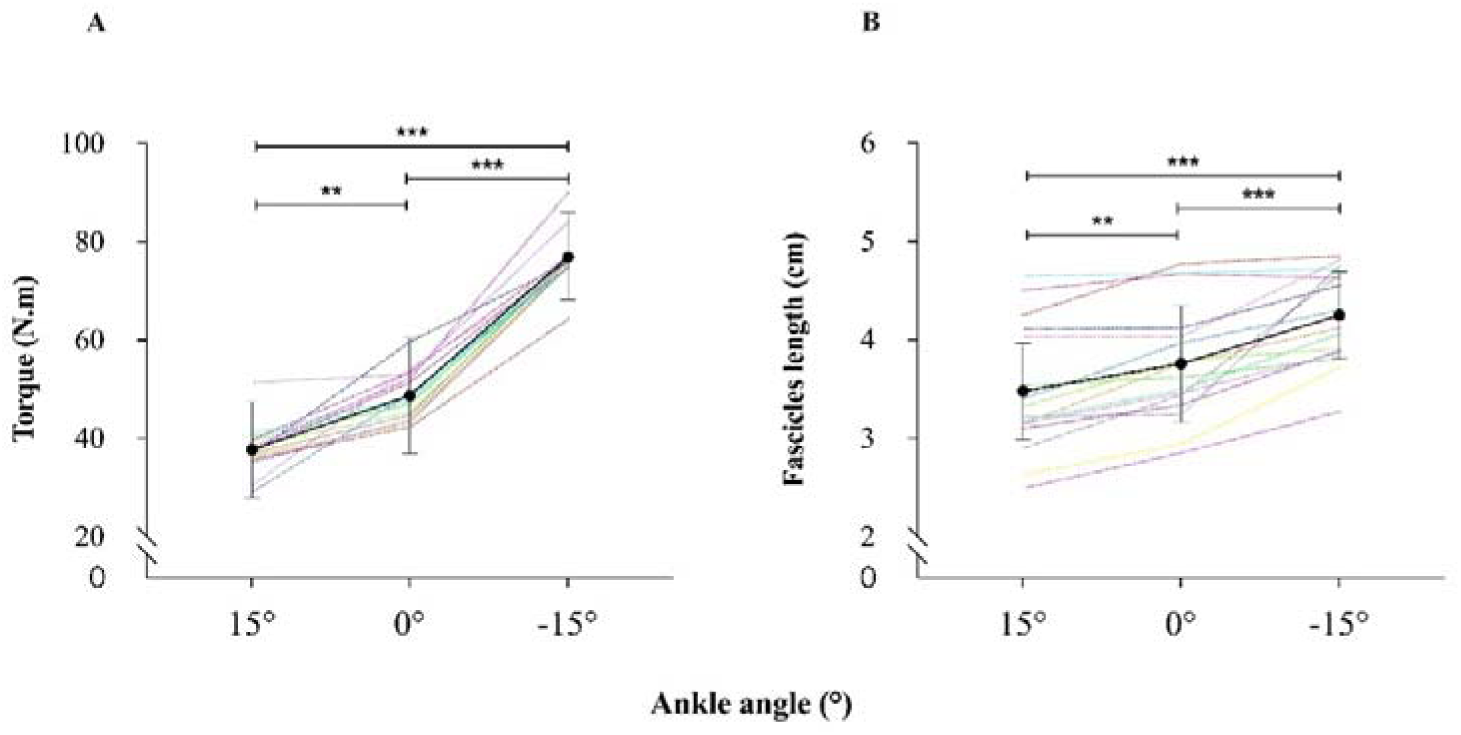
Changes in plantar flexors torques and SOL muscle length during 453 50% eccentric submaximal contraction according to ankle angle. Absolute and individual data (n = 16) are expressed as mean ± standard deviation A) Changes in plantar flexors torques according to ankle angle. ** Significant difference at P < 0.01 for the ankle angle effect. *** Significant difference at P < 0.001 for the ankle angle effect. B) Changes in soleus fascicles length according to ankle angle. ** Significant difference at P < 0.01 for the ankle angle effect. *** Significant difference at P < 0.001 for the ankle angle effect.

### Changes in fascicle length

The compiled images were used to track the length of the SOL fascicles using a customized MATLAB script that enabled tracking of linear tissues during eccentric submaximal contractions. The fascicle length was extracted at three different angles: +15° (stimulus point for short muscle length), 0° (stimulus point for intermediate muscle length) and -15° (stimulus point for long muscle length). One fascicle was tracked at each angle (Fig. 3). When the fascicles were not fully visible, those parts that were, as well as the deep aponeurosis, were extrapolated linearly to estimate the fascicle length.

### Statistical analysis

All descriptive statistics presented in the text, tables and figures are given as mean values ± standard deviation. The significance level for all analyses was set at P < 0.05. The normality of the data and homogeneity of variances were verified and validated using the Shapiro–Wilks W test and Levene test, respectively. Separate one-way repeated measures ANOVAs were used to assess changes in muscle torque and fascicle length at different ankle angles (15°, 0°, -15°) during eccentric submaximal contractions. Two-factor [contraction type (eccentric vs. isometric vs. concentric) × muscle length (short vs. intermediate vs. long)] repeated measures ANOVAs on contraction type and muscle length were used to compare SOL M_at_H_max_/M_max_, H_max_/M_max_, H_Slope_/M_Slope,_ H_1_/M_max_, V/M_max_, H_ref_/M_max_, H’/H_1_, H_D1_/H_test_ and H_Fac_/H_test_ among the conditions. Three-factor [contraction type (eccentric vs. isometric vs. concentric) × muscle length (short vs. intermediate vs. long) × experiment (experiment C vs. experiment D)] repeated measures ANOVAs on contraction type, muscle length and experiment were used to compare H_test_/M_max_. Three-factor [contraction type (eccentric vs. isometric vs. concentric) × muscle length (short vs. intermediate vs. long) × experiment (experiment A vs. experiment B vs. experiment C vs. experiment D)] repeated measures ANOVAs on contraction type, muscle length and experiment were used to compare SOL M_max_, EMG_RMS_/M_max_ for SOL and medial gastrocnemius, and tibialis anterior coactivation.

Analysis of covariance (ANCOVA) was performed to examine the effect of contraction type (eccentric vs. isometric vs. concentric) and muscle length (short vs. intermediate vs. long) on the amplitude of H’ with the V amplitude as a covariate. Whenever a significant main effect or interaction was detected, Tukey tests were performed for post hoc analysis. In addition, the relation between H_D1_/H_test_ and H_test_/M_max_ was examined using Pearson’s correlation coefficients. For this analysis, we used mean values per participant for each contraction type and each muscle.

The statistical analyses were performed using JASP (Version 0.17.2.1., Amsterdam, NED) and GraphPad Prism software (version 10.2.1; GraphPad Software Inc., San Diego, CA, USA).

## RESULTS

### Changes in passive and active torques

Torque during submaximal eccentric contraction (50% of maximal EMG_RMS_) was significantly different between each ankle angle (P < 0.001). It was 36.6% and 50.9% greater (P < 0.001) at an ankle angle of -15° than at 0° and +15°, respectively (Fig. 4A), and 22.5% higher (P < 0.001).

### Changes in fascicle length with ankle angle

Changes in fascicle length were calculated to ensure that the muscle lengths associated with angles of electrical nerve stimulation were different. Muscle lengths at -15° ankle angle were 12.9% and 22.1% (P < 0.001) greater than at 0° and 15° (Fig. 4B). The muscle length at 0° ankle angle was 8.2% (P = 0.011) greater than at 15°.

### EMG activity

We compared SOL and medial gastrocnemius EMG_RMS_/M_max_ between eccentric, isometric and concentric contractions at short, intermediate and long muscle lengths in order to detect those participants who produced the same agonist and synergistic muscle activations between all experimental conditions. No significant effects were found for either muscle (all P values > 0.90) (Table 1).

**Table 1.**
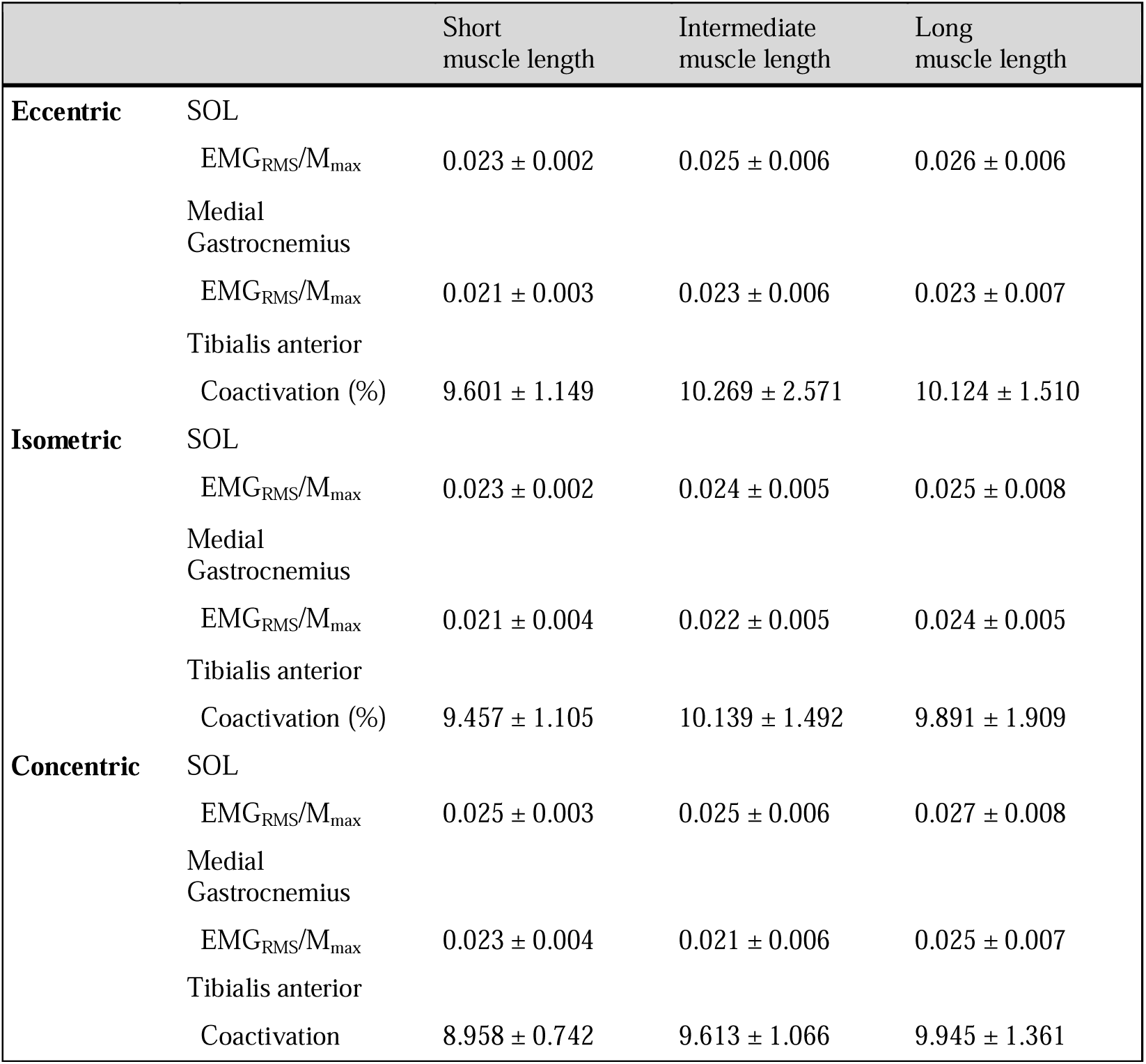
Effect of muscle length and contraction type on soleus, medial gastrocnemius and tibialis anterior EMG_RMS_ activity. Data are pooled for the four experiments and are expressed as mean ± standard deviation. EMG, electromyography; SOL, soleus; M_max_, maximal M wave; RMS, root mean square.

Tibialis anterior coactivation during plantarflexion was assessed to determine whether there was constant antagonist activation across all experimental conditions. No effect or interaction between conditions was found for the level of tibialis anterior coactivation (all P values > 0.081). The mean value of tibialis anterior coactivation was about 9.8% during all plantarflexor contractions (Table. 1). Antagonist activity, particularly reciprocal inhibition, may modulate the H reflex and consequently the recurrent inhibition pathway. Nevertheless, our findings indicate that while this phenomenon cannot be entirely discounted, it does not account for the modulation observed in our study.

### Effectiveness of activated Ia afferents to discharge a-motoneurons and reflex gain

Figure 5A illustrates raw traces showing the SOL H reflex and M wave evoked during eccentric, isometric and concentric contractions at short, intermediate and long muscle lengths in a representative participant. In this study, SOL H_max_/M_max_ and H_slope_/M_slope_ were assessed to estimate the effectiveness of activated Ia afferents to discharge _a_-motoneurons and the reflex gain, respectively.

**Figure 5.**
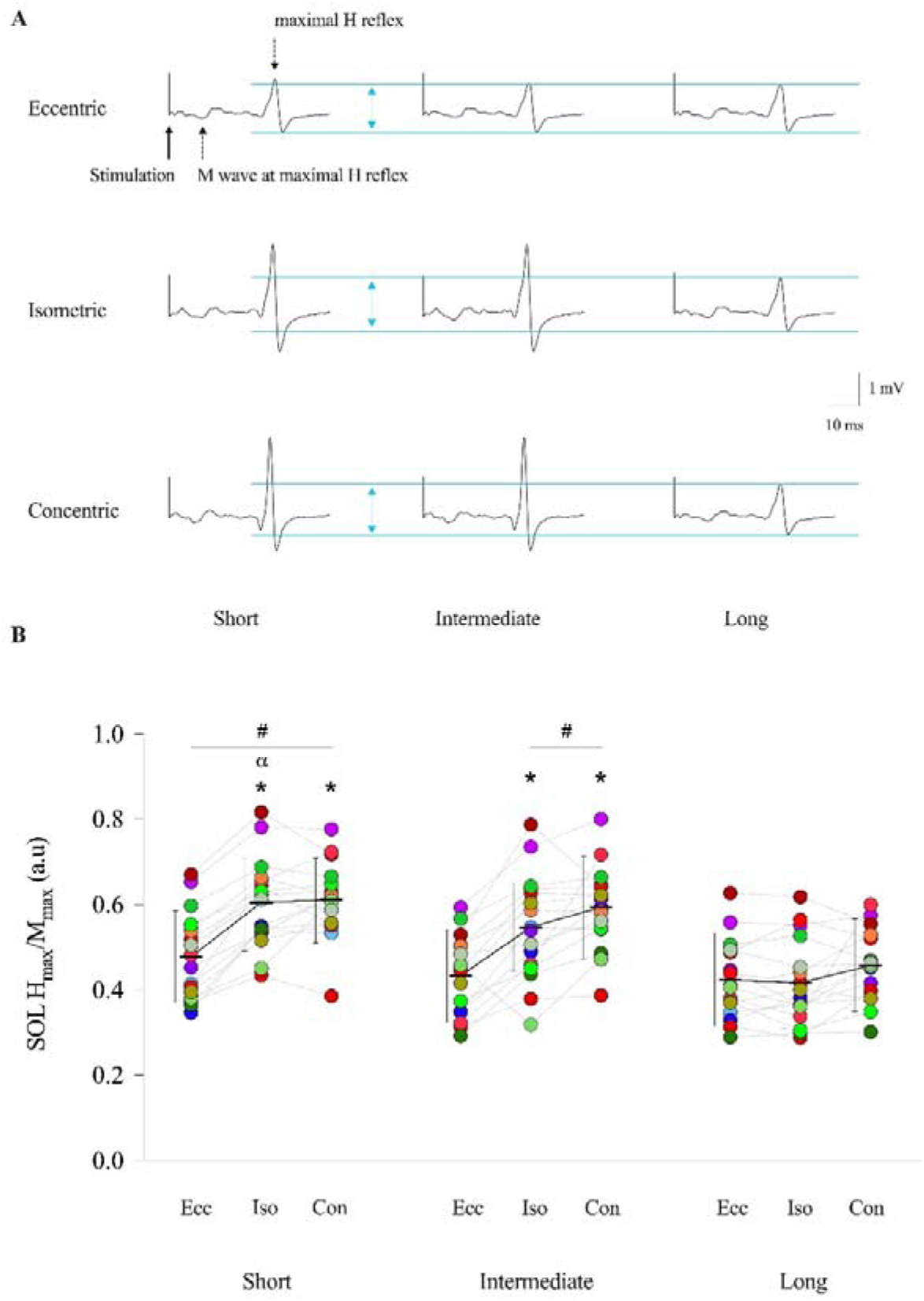
Change in maximal soleus H reflex normalized to the corresponding m 513 aximal M wave according to muscle length and contraction type. A) representative traces showing the soleus maximal H reflex evoked by posterior tibial nerve stimulation at short, intermediate and long muscle lengths during eccentric, isometric and concentric contractions. Blue dashed lines show the maximal H reflex amplitudes at long muscle length during each contraction type. B) Absolute and individual data (n = 16) are expressed as mean ± standard deviation. Changes in maximal soleus H reflex normalized to the corresponding maximal M wave at short, intermediate and long muscle lengths during eccentric, isometric and concentric contractions. Hmax, maximal soleus H reflex; Mmax, maximal M wave; Ecc, eccentric; Iso, isometric; Con, concentric. * Significantly different from eccentric contraction type. # Significantly different from long muscle length. α Significantly different from intermediate muscle length.

For SOL H_max_/M_max_, a contraction type × muscle length interaction effect (F _(4,60)_ = 20.679; P < 0.001; 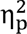 = 0.580) was found (Fig. 5B). First, muscle length influences the effect of contraction type on SOL H_max_/M_max_. At short muscle length, SOL H_max_/M_max_ was reduced by 26.5% and 29.3% during eccentric contraction compared with isometric and concentric contractions, respectively (P < 0.001). At intermediate muscle length, SOL H_max_/M_max_ was reduced by 23.1% and 32.9% during eccentric contraction compared with isometric and concentric contractions, respectively (P < 0.001). However, at long muscle length, no significant difference was found for SOL H_max_/M_max_ between eccentric contraction and the other contraction types (all P values > 0.671). No significant differences were found between isometric and concentric contraction types at any muscle length (all P values > 0.151).

Contraction type also influences the effect of muscle length on SOL H_max_/M_max_. During eccentric contraction, SOL H_max_/M_max_ was only reduced by 12.1% at long muscle length compared with short muscle length (P = 0.022), and no difference was found between long and intermediate or intermediate and short muscle lengths (all P values > 0.107). During isometric contraction, SOL H_max_/M_max_ was reduced by 31.6% and 44.9% at long muscle length compared with intermediate and short muscle lengths, respectively (P < 0.001). In addition, SOL H_max_/M_max_ was reduced by 10.1% at intermediate compared with short muscle length (P = 0.012). During concentric contraction SOL H_max_/M_max_ was reduced by 31.3% and 36.8% at long muscle length compared with intermediate and short muscle lengths (P < 0.001), but no significant differences were found between intermediate and short muscle lengths (P = 1.000).

For the SOL H_slope_/M_slope_, a significant contraction type × muscle length interaction (F = 3.751; P = 0.009; 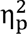 = 0.200) was found (Fig. 6). Primarily, muscle length influences the effect of contraction type on H_slope_/M_slope_.

**Figure 6.**
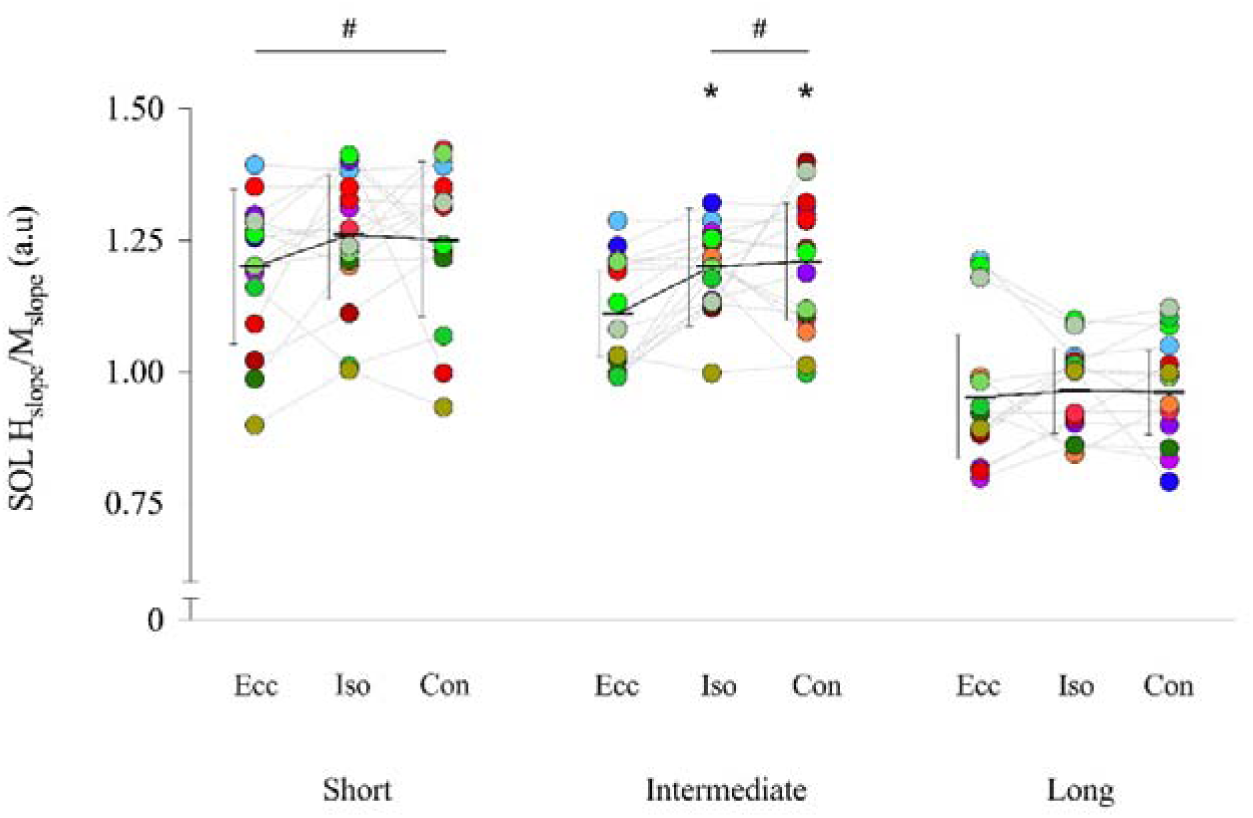
Changes in soleus H reflex slope development and M wave slope development ratio according to muscle length and contraction type. Absolute and individual data (n=16) are expressed as mean ± standard deviation. H_slope_; soleus H reflex slope development; M_slope_, M wave slope development; Ecc, eccentric; Iso, isometric; Con, concentric. * Significantly different from eccentric contraction type. # Significantly different from long muscle length.

At short muscle length, no significant difference was found for SOL H_slope_/M_slope_ during eccentric contraction compared with isometric and concentric contraction types (all P values > 0.688). At intermediate muscle length, SOL H_slope_/M_slope_ was reduced by 9.6% and 12.3% during eccentric contraction compared with isometric and concentric contractions, respectively (P = 0.002 and P = 0.033). However, at long muscle length no significant difference was found for SOL H_slope_/M_slope_ between eccentric contraction and the other contraction types (all P values = 1.000). No significant differences were found between isometric and concentric contraction types for any muscle length (all P values = 1.000).

Contraction type also influences the effect of muscle length on SOL H_slope_/M_slope_. During eccentric contraction, SOL H_slope_/M_slope_ was found to be reduced by 18.5% at long muscle length compared with short muscle length (P < 0.001), and there was no difference between long and intermediate muscle lengths (P = 0.143). During isometric contraction, SOL H_slope_/M_slope_ was reduced by 23.7% and 27.5% at long muscle length compared with intermediate and short muscle length (P < 0.001). During concentric contraction SOL H_slope_/M_slope_ was reduced by 27.2% and 25.5% at long muscle length compared with intermediate and short muscle lengths (P < 0.001). No significant differences were found between intermediate and short muscle lengths for any contraction type (all P values = 1.000).

We measured SOL M_max_ to assess the stability of EMG recording across the different experimental conditions. For M_max_, no significant effect or interaction between conditions was found (all P values > 0.201, Table 2).

**Table 2.**
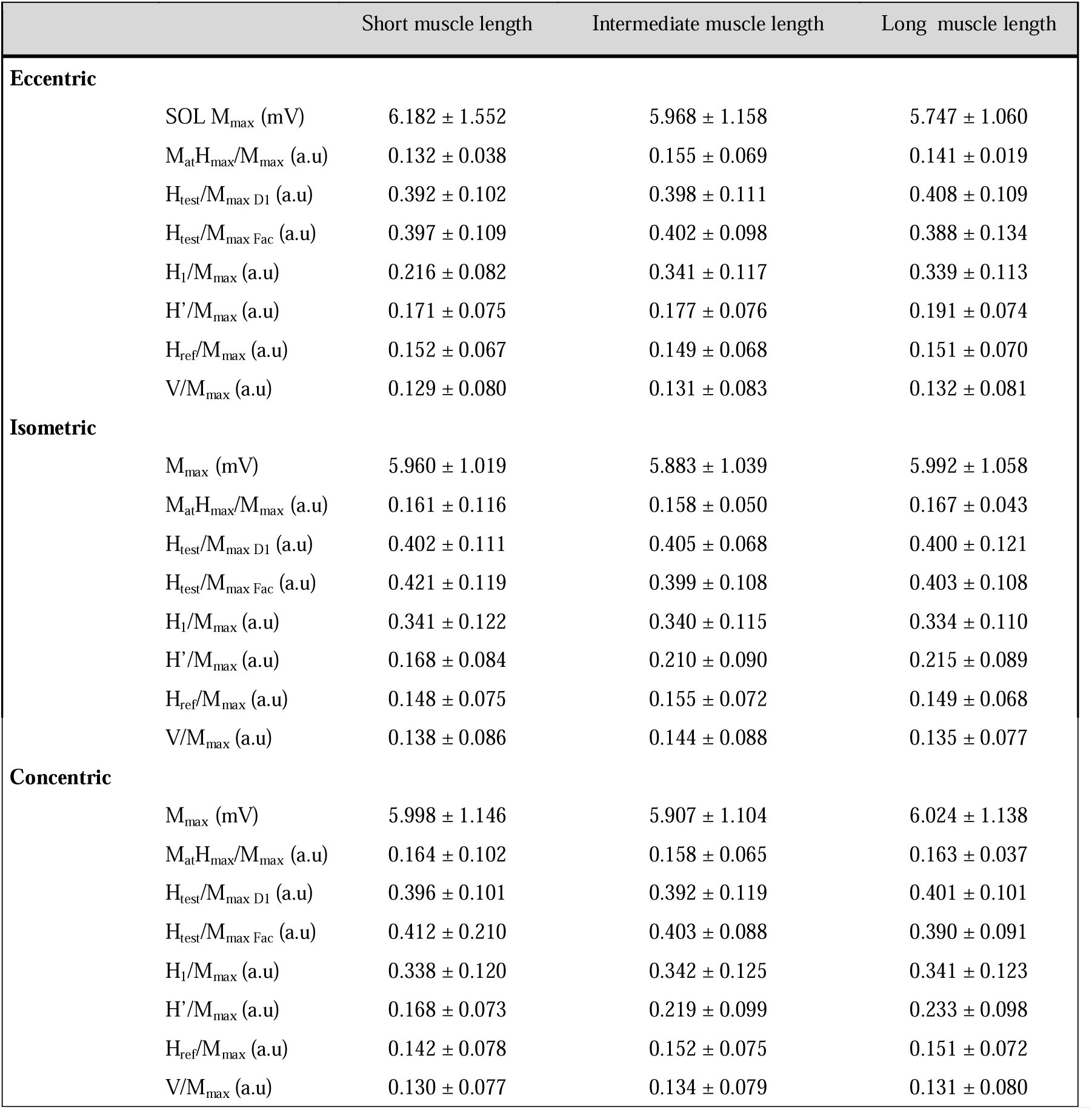
Effect of contraction type and muscle length on soleus evoked potentials. Absolute data (n = 16 and n = 12 for heteronymous Ia facilitation) are expressed as mean ± standard deviation. SOL, soleus; M_max_, maximal M wave; M_at_H_max_, M wave corresponding to maximal soleus H reflex; H_test_, non-conditioned soleus H reflex corresponding to its test value for D1 and heteronymous Ia facilitation methods; H_1_, conditioning H reflex amplitude for paired H reflex technique; H’, test H reflex for paired H reflex technique; H_ref_, reference H reflex paired H reflex technique; V, V wave.

We used SOL M_at_H_max_/M_max_ to determine whether the same numbers of both efferent and potentially afferent nerve fibres were effectively stimulated under the different conditions (Schieppati 1987). Our findings indicate that the amplitude of M_max_ and M_at_H_max_ remain consistent, suggesting that an equivalent proportion of _a_-motoneurons is activated across all conditions (Table 2). Thus, the observed H_max_/M_max_ modulations cannot be attributed to recording conditions but rather to neural mechanisms (Schieppati, 1987; Duclay & Martin, 2005).

### Recurrent inhibition

Figure 7A illustrates raw traces showing the conditioning (H_1_) and test (H’) H reflex measured during eccentric, concentric and isometric contractions at long, intermediate and short muscle lengths in a representative participant. In this study, SOL H’/H_1_ was assessed to estimate the level of recurrent inhibition.

**Figure 7.**
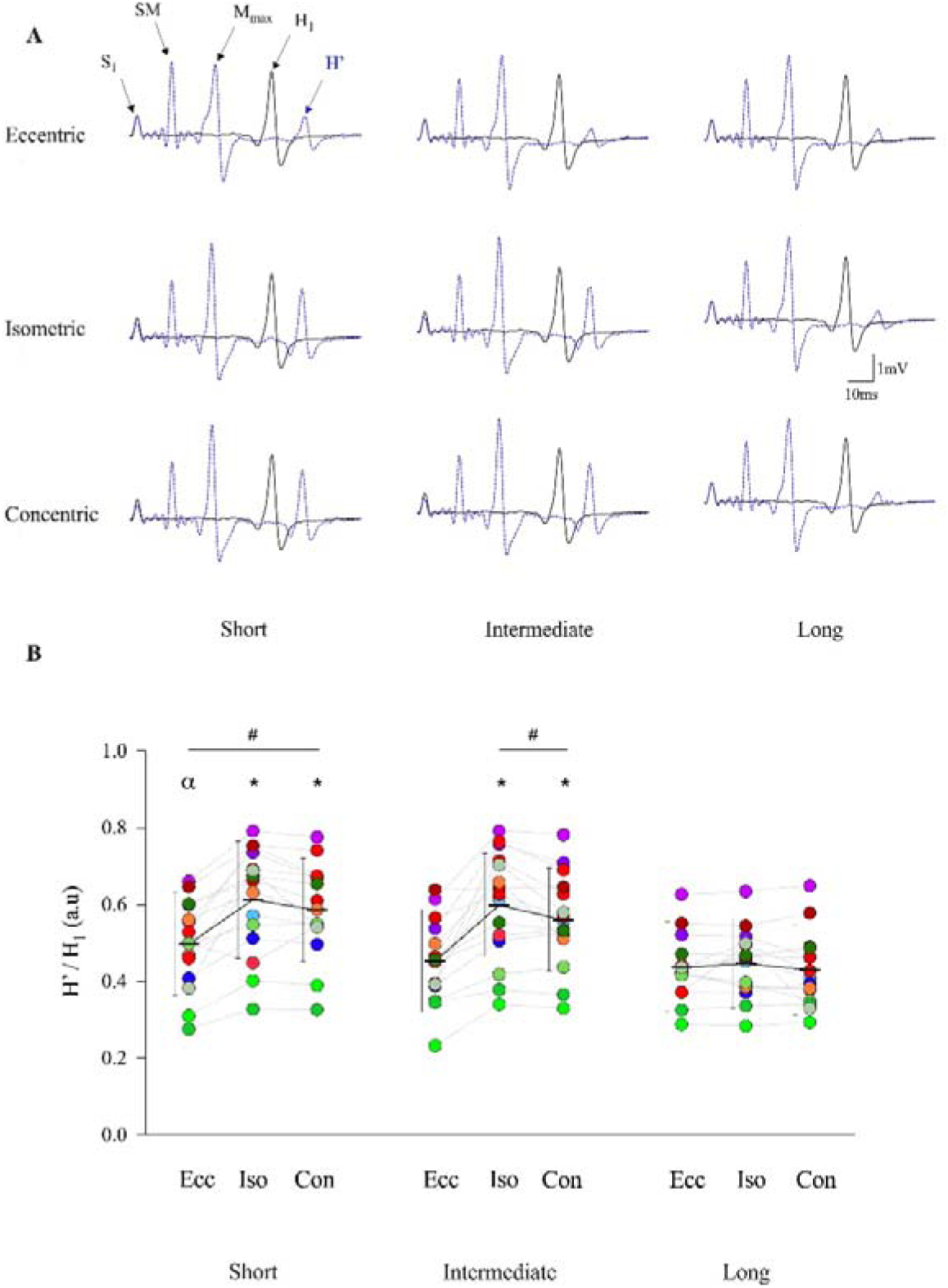
Change in recurrent inhibition according to muscle length and contraction type. A) Representative traces showing conditioning soleus H reflex (H_1_) and test soleus H reflex (H’). The test H reflex (H’) was elicited through S_1_ + supramaximal intensity stimulations on the posterior tibial nerve, with a 10-ms time interval, during eccentric, concentric and isometric contractions at long, intermediate and short muscle lengths. Black line corresponds to the conditioning H reflex (H_1_) and blue line to the test H reflex (H’). S_1_, stimulation inducing H_1_; supramaximal intensity, stimulation evoking M_max_; M_max_, maximal M wave; H_1_, conditioning H reflex evoked by S_1_; H’, test H reflex evoked by S_1_ + supramaximal intensity. B) Absolute and individual data (n = 16) are expressed as mean ± standard deviation. H’ and H_1_ are expressed as a ratio. The electrical intensity to evoke H_1_ was adjusted in each muscle length during all contractions to ensure that the conditioning H reflex was present without a concomitant M wave. H’, test soleus H reflex; H_1_, conditioning soleus H reflex_;_ Ecc, eccentric; Iso, isometric; Con, concentric. * Significantly different from eccentric contraction type. # Significantly different from long muscle length. a Significantly different from intermediate muscle length.

For SOL H’/H_1_, a significant contraction type × muscle length interaction effect (F_(4,60)_ = 18.628; P < 0.001; 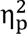 = 0.554) was found (Fig. 7B).

First, muscle length influenced the effect of contraction type on SOL H’/H_1_. At short muscle length, SOL H’/H_1_ was reduced by 23.8% and 18.1% during eccentric contraction compared with isometric and concentric contractions, respectively (P < 0.001). At intermediate muscle length SOL H’/H_1_ was reduced by 34.5% and 24.9% during eccentric contraction compared with isometric and concentric contractions, respectively (P < 0.001). However, at long muscle length, no significant difference was found for SOL H’/H_1_ between eccentric contractions and the other contraction types (all P values = 1.000). No significant differences were found between isometric and concentric contraction types at any muscle length (all P values > 0.218).

Contraction type also influences the effect of muscle length on SOL H’/H_1_. During eccentric contractions, SOL H’/H_1_ was only reduced by 15.8% at long muscle length compared with short muscle length (P = 0.002), and no difference was found between long and intermediate muscle lengths (P = 1.000). However, SOL H’/H_1_ was reduced by 10.4% at intermediate compared with short muscle lengths (P = 0.047). During isometric contractions, SOL H’/H_1_ was reduced by 33.6% and 37.4% at long muscle length compared with intermediate and short muscle lengths (P < 0.001), but no significant difference was found between intermediate and short muscle lengths (P = 1.000). During concentric contractions, SOL H’/H_1_ was reduced by 28.3% and 35.5% at long muscle length compared with intermediate and short muscle length (P < 0.001), but no significant difference was found between intermediate and short muscle lengths (P = 0.706).

SOL H_1_/M_max_ was assessed to ensure that the number of motoneurons involved in the H_1_ conditioning discharge and potentially affected by recurrent inhibition could be considered similar across all experimental conditions. No significant effects or interactions were found for SOL H_1_/M_max_ (P = 0.500, Table 2). Our results indicate that the number of motoneurons involved in the H_1_ conditioning discharge and potentially affected by recurrent inhibition can be considered similar between contraction types and muscle length.

Our work produced mean H_1_/M_max_ and H’/M_max_ values of 0.334 ± 0.011 and 0.194 ± 0.009, respectively, regardless of contraction type, and 0.348 ± 0.016 and 0.195 ± 0.013, respectively, regardless of muscle length. Comparison with previous literature (Katz & Pierrot-Deseilligny, 1999) appears to confirm that the intensity of our S_1_ stimulus elicits H_1_ and H’ responses corresponding to the descending phase of the H’/H_1_ curve, which limits contamination of H’ by the post-spike afterhyperpolarization mechanism.

SOL H_ref_/M_max_ was calculated to check whether the modulations of H’ related to muscle length and contraction type were greater than those of H_ref_. No significant effects or interactions were found for SOL H_ref_/M_max_ across the experimental conditions studied (all P values > 0.476, Table 2). Given that SOL H_ref_/M_max_ and H_1_/M_max_ remain constant while the ratio H’/H_1_ varies significantly, the observed variation in H’ exceeds that of H_ref_. This indicates that the variations in H’ are likely due to changes in the amount of recurrent inhibition.

V/M_max_ was used to determine whether the contamination of H’ by V remained consistent across all conditions. A repeated measures ANOVA revealed no significant effects or interactions for V/M_max_ among test conditions (all p-values > 0.264, see Table 2). These results confirm that V/M_max_ does not account for the variations observed in H’.

A repeated measures ANCOVA was conducted to ensure that the effects of contraction type and muscle length on H′ were not confounded by V wave (covariate) contamination. Significant effects were found for the covariate (F_(1,134)_ = 12.174; P < 0.001; 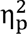 = 0.083) and muscle length (F_(2,134)_ = 3.693; P = 0.027; 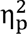 = 0.052), but no interaction between muscle length and contraction type was observed (F_(4,134)_ = 0.451; P = 0.771;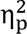 = 0.013). Post hoc analyses revealed that, even after checking for the effect of the covariate V, H′ was significantly reduced at long compared with short muscle lengths (P = 0.028).

Furthermore, these results demonstrate that the contamination of H′ by V was consistent across all contraction types, confirming the findings of Barrué-Bélou et al. (2018, 2019), which indicate that this contamination does not depend on muscle length. These results suggest that the effects of neither muscle contraction type nor muscle length on H′/H_1_ ratio can be attributed to the contamination of H′ by V.

### Post-activation by PAD (D1 method)

Figure 8A illustrates raw traces showing the conditioned (H_D1_) and non-conditioned (H_test_) H reflex measured during eccentric, concentric and isometric contractions at long, intermediate and short muscle lengths in a representative participant. SOL H_D1_/H_test_ was assessed to estimate the level of post-activation depression by PAD.

**Figure 8.**
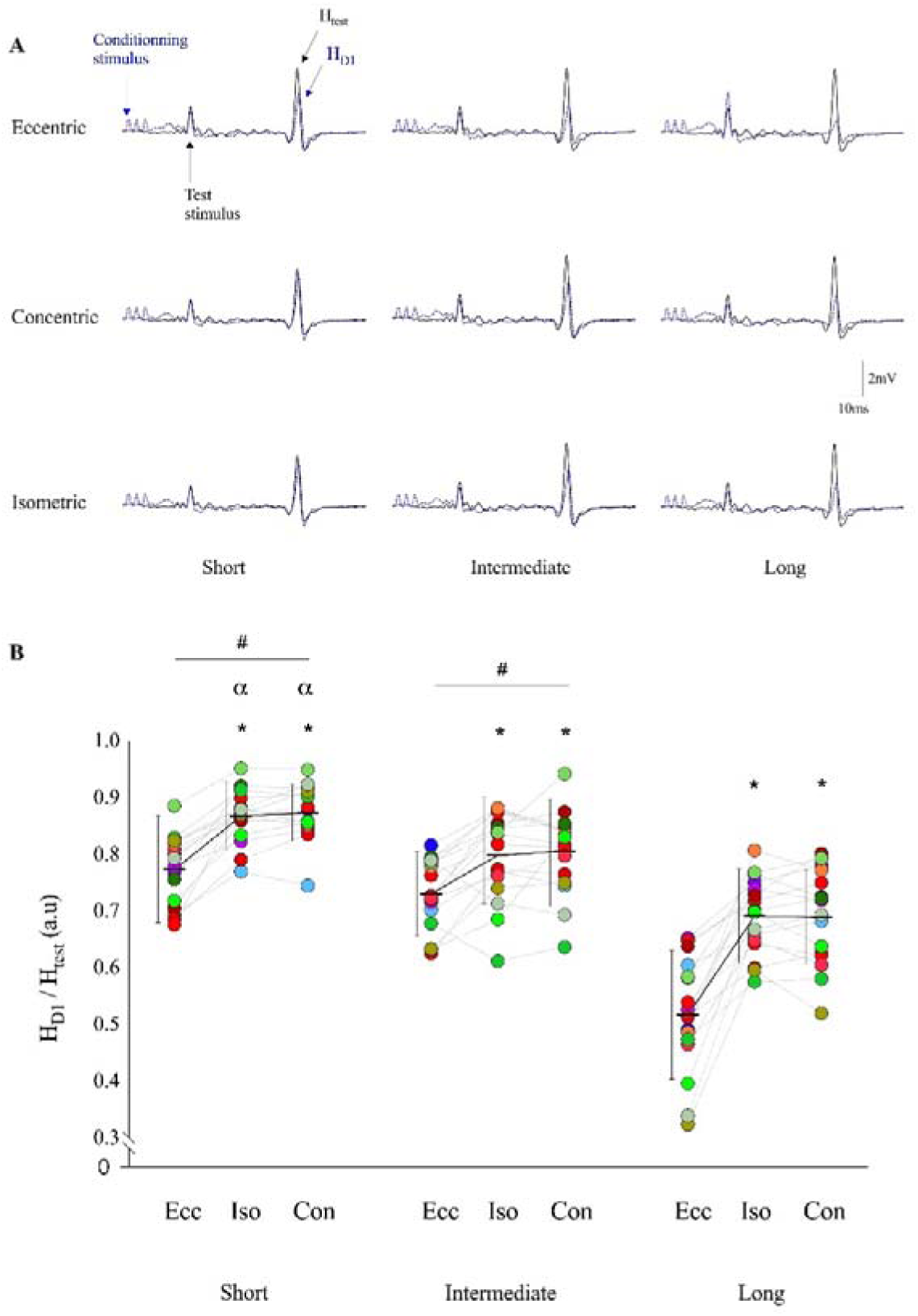
Changes in post-activation depression by primary afferent depolarization according to muscle length and contraction type. A) Representative traces showing test H reflex (H_test_) and the conditioned soleus H reflex (H_D1_) evoked by fibular nerve stimulation preceding the posterior tibial nerve stimulation at a conditioning test interval of 21 ms during eccentric, concentric and isometric contractions at long, intermediate and short muscle lengths. Black lines correspond to the non-conditioned H reflex (H_test_) and blue dashed lines to the conditioned H reflex (H_D1_). B) Absolute and individual data (n =16) are expressed as mean ± standard deviation. H_D1_ and H_test_ are expressed as a ratio during eccentric, isometric and concentric contractions at short, intermediate and long muscle lengths. The electrical intensity to evoke H_test_ was normalized during all contractions at each muscle length. H_D1_, conditioned soleus H reflex; H_test_, non-conditioned soleus H reflex; Ecc, eccentric; Con, concentric; Iso, isometric. * Significantly different from eccentric contraction type. # Significantly different from long muscle length. a Significantly different from intermediate muscle length.

For SOL H_D1_/H_test_ a significant interaction of contraction type × muscle length (F _(4,60)_ = 6.914; P < 0.001; 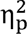 = 0.316) was found (Fig. 8B).

First, muscle length influences the effect of contraction type on SOL H_D1_/H_test_. At short muscle length, SOL H_D1_/H_test_ was reduced by 12.2% and 13.1% during eccentric contraction compared with isometric and concentric contractions, respectively (P = 0.001). At an intermediate muscle length, SOL H_D1_/H_test_ was reduced by 9.3% and 10.4% during eccentric contraction compared with isometric and concentric contractions, respectively (P < 0.001). As illustrated in Figure 8B, the reductions were even greater at a long muscle length. SOL H_D1_/H_test_ was reduced by 33.9% and 33.5% during eccentric contraction compared with isometric and concentric contractions, respectively (P < 0.001). No significant differences were found between isometric and concentric contraction types at any muscle length (all P values =1.000).

Contraction type also influences the effect of muscle length on SOL H_D1_/H_test_. During eccentric contraction, SOL H_D1_/H_test_ was reduced by 41.3% and 49.7% at long muscle length compared with intermediate and short muscle length, respectively (P < 0.001). No significant difference was found between intermediate and short muscle lengths (P = 0.417). During isometric contraction SOL H_D1_/H_test_ was reduced by 15.4% and 25.4% at long muscle length compared with intermediate and short muscle length, respectively (P < 0.001). In addition, SOL H_D1_/H_test_ was reduced by 8.7% at intermediate compared with short muscle length (P = 0.024). During concentric contraction, SOL H_D1_/H_test_ was reduced by 16.9% and 26.7% at long muscle length compared with intermediate and short muscle length, respectively (P < 0.001). SOL H_D1_/H_test_ was reduced by 8.4% at intermediate compared with short muscle length (P = 0.029).

SOL H_test_/M_max_ was measured to assess the stability of the H reflex test between conditions during post-activation depression by PAD assessment (Table 2). No significant effects or interactions were observed for SOL H_test_/M_max_ (all P values > 0.167). In our investigation of post-activation depression by PAD and heteronymous Ia facilitation, we adjusted the stimulus intensity to maintain a constant H_test_/M_max_, ensuring consistent amplitude of the non-conditioned H reflex across experimental conditions. However, this adaptation of stimulus intensity means that a different number of Ia afferents can contribute to the reflex under different conditions. Despite this, correlation analysis revealed no significant relationship between H_D1_/H_test_ and H_test_/M_max_ results (r = 0.172; P = 0.523). These findings are consistent with previous results reported by Papitsa et al. (2022). Therefore, the observed modulations in H_D1_ and H_Fac_ amplitudes are likely attributable to variations in excitatory and/or inhibitory inputs at the presynaptic level, as supported by earlier studies (Baudry & Enoka, 2009; Baudry & Duchateau, 2012).

### Heteronymous Ia facilitation (HF)

Figure 9A illustrates raw traces showing the conditioned (H_Fac_) and non-conditioned (H_test_) H reflex measured during eccentric, isometric and concentric contractions at short, intermediate and long muscle lengths in a representative participant.

**Figure 9.**
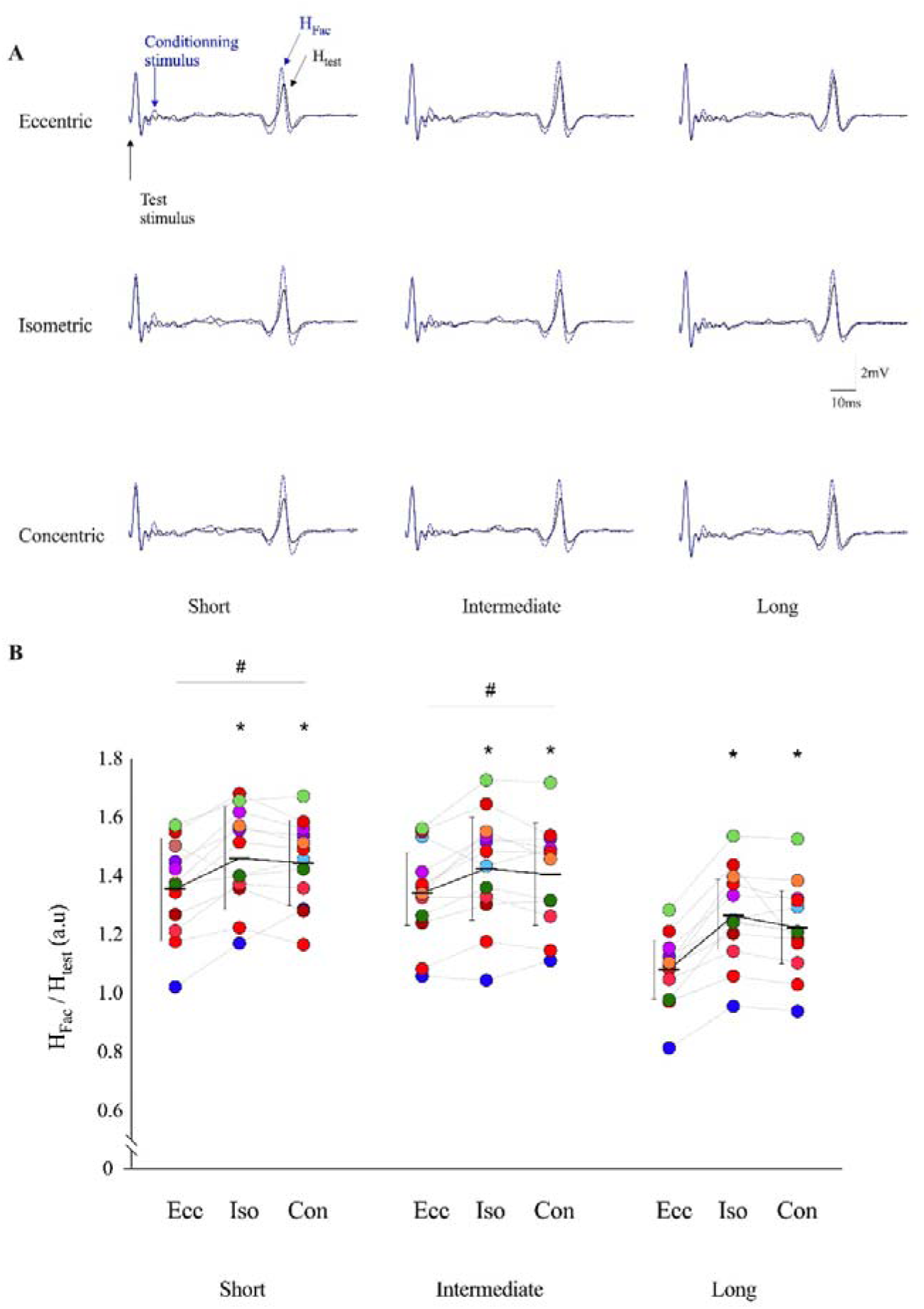
Changes in heteronymous Ia facilitation according to muscle length and contraction type. A) Representative traces showing the non-conditioned soleus H reflex (H_test_) and conditioned H reflex (H_Fac_). H_test_ and H_Fac_ were evoked by posterior tibial nerve stimulation and the conditioned soleus H reflex (H_Fac_) evoked by femoral nerve stimulation following the posterior tibial nerve stimulation at a conditioning test interval ranging from 9 to 1 ms during eccentric, isometric and concentric contractions at short, intermediate and long muscle lengths. Black lines correspond to the non-conditioned H reflex (H_test_) and blue dashed lines to the conditioned H reflex (H_Fac_). B) Absolute and individual data (n =12) are expressed as mean ± standard deviation. H_Fac_ and H_test_ are expressed as a ratio during eccentric, concentric and isometric contractions at long, intermediate and short muscle lengths. The electrical intensity to evoke H_test_ was normalized during all contractions at each muscle length. H_Fac_, conditioned soleus H reflex; H_test_, non-conditioned soleus H reflex; Ecc, eccentric; Con, concentric; Iso, isometric. * Significantly different from eccentric contraction type. # Significantly different from long muscle length.

For SOL H_Fac_/H_test_, a significant contraction type × muscle length interaction (F _(4,44)_ = 5.184; P < 0.002; 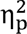 = 0.320) was found (Fig. 9B).

First, muscle length influences the effect of contraction type on SOL H_Fac_/H_test_. At a short muscle length, SOL H_Fac_/H_test_ was reduced by 7.7% and 6.5 % during eccentric contraction compared with isometric and concentric contractions, respectively (P = 0.001 and P = 0.004). At an intermediate muscle length, SOL H_Fac_/H_test_ was reduced by 6.2% during eccentric contraction compared with isometric contractions (P = 0.008). As illustrated in Figure 9B, the reductions were even greater at a long muscle length. SOL H_Fac_/H_test_ was reduced by 17.1% and 13.3% during eccentric contraction compared with isometric and concentric contractions, respectively (P < 0.001). No significant differences were found between isometric and concentric contraction types at any muscle length (all P values > 0.472).

Contraction type also influences the effect of muscle length on SOL H_Fac_/H_test_. During eccentric contraction, we found that SOL H_Fac_/H_test_ was reduced by 24.2% and 25.5% at long muscle length compared with intermediate and short muscle lengths, respectively (P < 0.001). During isometric contraction SOL H_Fac_/H_test_ was reduced by 12.7% and 15.5% at long muscle compared with intermediate and short muscle lengths, respectively (P < 0.001). During concentric contraction SOL H_Fac_/H_test_ was reduced by 14.7% and 18.1% at long muscle length compared with intermediate and short muscle lengths (P < 0.001). No significant differences were found between intermediate and short muscle lengths for any contraction type (all P values > 0.460).

## DISCUSSION

The objective of this study was to examine the impact of muscle length on the effectiveness of activated Ia afferents to discharge α-motoneurons, as well as reflex gain, recurrent inhibitory, post-activation depression by PAD and heteronomous Ia facilitation during eccentric, isometric and concentric contraction types. Our findings offer valuable new insights into the influence of muscle length and contraction type on the regulation of Ia afferent inputs to motoneuron excitability. They demonstrate a reduction in the effectiveness of activated Ia afferents to discharge α-motoneurons and reflex gain during eccentric contractions at intermediate and short muscle lengths. In contrast, for long muscle length, the effectiveness of activated Ia afferents to discharge α-motoneurons was similar for all contraction types. Interestingly, this specific regulation seems to stem from changes in the extent of recurrent inhibition. Overall, our study demonstrates a specific interaction between the effects of muscle length and contraction type on inhibitory mechanisms acting on the SOL muscle at the spinal level.

### The effect of contraction type on spinal regulatory mechanisms is influenced by changes in muscle length

This study is the first to examine the influence of muscle length on the effectiveness of activated Ia afferents to discharge _a_-motoneurons during isometric and dynamic muscle contractions. Furthermore, we used a dual approach (i.e., H_max_ and H_slope_) to concomitantly investigate the underlying regulatory mechanisms.

Our original results indicate that the typical reductions in H_max_/M_max_ and H_slope_/M_slope_ ratios observed during eccentric contractions at intermediate muscle lengths, compared with isometric and concentric contractions, is not observed at longer muscle lengths. This suggests that variations in the effectiveness of activated Ia afferents to discharge _a_-motoneurons during eccentric contractions at longer muscle lengths can be explained by changes in reflex gain. Reflex gain, which we measured in all test conditions using recruitment curves with constant increases in stimulation intensity, is an indicator of the sensitivity of motoneurons to afferent input, specifically indicating how changes in afferent input influence the rate of motoneuron discharge (Funase *et al*., 1994). Reflex gain was measured using recruitment curves with constant increases in stimulation intensity in all conditions. This method is commonly used in static conditions as proposed by (Hwang, 2002). Therefore, the similar modulations of the effectiveness of activated Ia afferents to discharge _a_-motoneurons and reflex gain during eccentric contractions at long muscle lengths confirms that these specific modulations result from changes in the regulation of the balance between inhibition and excitation acting at the spinal level (Funase *et al*., 1994).

Our results show that the increase in post-activation depression by PAD during eccentric contractions occurs for short (mean 12.6%), intermediate (mean 9.8%) and long muscle lengths, with the greatest increase observed at long muscle lengths (mean 33.7%). In this latter condition, we could have expected that a reduction in reflex gain occurs, which would indicate a decrease in the effectiveness of afferent discharges reaching _a_-motoneurons. Surprisingly, our results showed similar levels of reflex gain at long muscle length for all contraction types. This unexpected result suggests that changes in _a_-motoneuron excitability, and therefore in post-synaptic mechanisms, may play an important role in maintaining reflex gain despite increased post-activation depression (presynaptic mechanism) during eccentric contractions at long muscle length.

Our mechanical data indicate that the torque generated during eccentric contractions at long muscle lengths is greater than that produced at intermediate or short muscle lengths. It can thus be postulated that an increase in intra- and extra-fusal muscle tension enhances afferent discharges (Ia and II), which may elevate levels of post-activation depression (Colard *et al*., 2023) during eccentric contractions at long muscle lengths. It is well documented, notably by Burke *et al*. (1978), that neuromuscular spindle firing rates are significantly faster during eccentric contractions than during concentric ones. However, it must be considered that this effect could be counterbalanced by the regulatory influence of gamma motoneurons on intrafusal fibres, maintaining a balanced discharge reaching the afferent Ia terminal (Petersen *et al*., 2007). Thus, previous studies have privileged an eccentric-specific supraspinal influence that would regulate the activity of post-activation depression by PAD (Grosprêtre *et al*., 2014; Colard *et al*., 2023). It is plausible that specific changes in supraspinal (i.e., eccentric neural specificity) control mechanisms could potentially affect GABAergic interneuron threshold excitability (Colard *et al*., 2023). Furthermore, the additional afferent discharge due to the long muscle length could generate greater than usual activation of GABA_A_ receptors. This increased activation of GABA_A_ receptors could produce greater PAD with larger facilitating spikes (Metz *et al*., 2023*a*). Thus the additional early and late post-activation depression could be explained by an amplification of the following mechanisms: (1) decreased afferent excitability due to collision of orthodromic action potentials with antidromic PAD-evoked spikes; (2) transmitter depletion at the Ia terminal by PAD-evoked spikes and/or (3) inhibition of Ia afferent terminals by GABA_B_ receptors indirectly mediating presynaptic inhibition (Hari *et al*., 2022; Metz *et al*., 2023*a*). In order to confirm the involvement of these mechanisms, we used the heteronymous Ia facilitation.

The complementary between the D1 method (decrease in H_D1_/H_test_) and heteronymous Ia facilitation (decrease in H_Fac_/H_test_) is based on the hypothesis that a constant conditioning stimulus would consistently excite GABAergic interneurons (D1 method) or motoneurons (HF), resulting in stable post-activation by PAD activity and additional stable heteronymous Ia facilitation, unless the post-activation by PAD changes (Baudry & Enoka, 2009). Therefore, our results demonstrate an increase in post-activation depression by PAD and a concomitant reduction in heteronymous Ia facilitation during eccentric contractions at long muscle length. Consequently, our measurements do reveal changes in GABAergic interneuron activity.

Nevertheless, the stability of reflex gain and the effectiveness of activated Ia afferents to discharge _a_-motoneurons despite an increase in inhibition upstream of the synapse during eccentric contraction at long muscle length underlines the necessity for a more comprehensive analysis of post-synaptic mechanisms, particularly recurrent inhibition whose role has already been observed when comparing contraction types.

To our knowledge, the present study is the first to investigate the effect of muscle length on recurrent inhibition during different contraction types. Our data corroborates the findings of previous studies and confirm that, at intermediate lengths, the H’/H_1_ ratio is reduced during eccentric contractions compared with isometric and concentric contractions (Barrué-Belou *et al*., 2018; Papitsa *et al*., 2022). We found that these results can be extended to short muscle lengths, which also show a reduced H’/H_1_ during eccentric contractions relative to isometric or concentric ones. However, we observed that the H’/H_1_ ratio during eccentric contractions performed at long muscle lengths is similar to that observed during isometric and concentric contractions. This finding suggests that recurrent inhibition during eccentric contractions may be modulated differently by long muscle lengths compared with intermediate and short muscle lengths. As previously indicated, post-activation depression by PAD is intensified during eccentric contractions of long muscle length. It is probable that this inhibition is the result of two associated factors: a reduction in the motor threshold of GABAergic interneurons by supraspinal control, i.e., specific to eccentric contractions (Grosprêtre *et al*., 2014; Papitsa *et al*., 2022; Colard *et al*., 2023), and increased activity of Ia and II afferent discharges associated with greater muscle lengths (i.e. static length variation). In such extreme stretching conditions, excessive inhibition of peripheral pathways could lead to over-inhibition of motoneuron activity, limiting motor output and failing to produce the required force. We suggest that some of the additional afferent discharge due to increased muscle length may facilitate type II afferent discharges, which could simultaneously reach Renshaw cells via an inhibitory interneuron (Jankowska, 1992), reducing their activity to prevent deficits in α-motoneuron activation and ensure adequate motor output (Windhorst, 1996). This mechanism could serve as a corrective response to perturbations, enabling the nervous system to adapt to high mechanical stress. This hypothesis is supported by the fact that type II afferents have an excitability threshold double that of type I afferents (Simonetta-Moreau *et al*., 1999). Consequently, this phenomenon of facilitation of type II afferents over Renshaw inhibition only occurs at long lengths where tension on neuromuscular spindle is maximal compared with short lengths. It is acknowledged that other concurrent mechanisms may be involved, such as alterations in the discharge of recurrent collaterals (Katz & Pierrot-Deseilligny, 1999) or an increased discharge of synergistic activity (Meunier *et al*., 1994). However, our results indicate comparable activity of the medial gastrocnemius muscle between the different conditions, suggesting that the latter possibility is limited.

### Effect of muscle length on spinal regulatory mechanisms

One of the original findings of this study was that SOL H_max_/M_max_ and H_slope_/M_slope_ were reduced at long muscle length compared with short muscle length for all contraction types. Our results demonstrate that this modulation is explained by a greater post-activation by PAD and recurrent inhibition acting at the spinal level. However, it could be pertinent to suggest that excitatory inputs from lumbar propriospinal neurons to α-motoneurons may be reduced. For example, a decrease in the sensitivity of these lumbar propriospinal neurons to II afferent discharge from (i) PAD activity, (ii) noradrenergic inputs from the locus coeruleus (Jankowska, 1992; Marchand-Pauvert *et al*., 2005) and (iii) extrapyramidal pathways could reduce the effectiveness of activated Ia afferents to discharge α-motoneurons (Jankowska & Edgley, 2010). Although cutaneous afferents can play an important role in the observed modulations, their involvement seems limited. Indeed, these afferents could potentially reduce post-activation depression by PAD via an inhibitory interneuron, which would modulate the activity of glutamatergic interneurons (Rudomin & Schmidt, 1999). However, the results of our study indicate an increase in post-activation depression by PAD. This result suggests that cutaneous afferents do not have a significant effect in this specific context.

We have demonstrated original findings indicating that the regulation of post-activation depression by PAD is greater during voluntary contraction at long compared with intermediate and short muscle lengths. Our mechanical assessments reveal a notable increase in active torque (i.e., an increase in intra- and extra-fusal muscle tension) at long muscle length, compared with intermediate or short muscle lengths. This high tension applied on neuromuscular spindle at long muscle length could generate a very large afferent influx towards the Ia terminal, which could generate reflex activity on the agonist muscle, thus altering the quality of movement. To avoid excessive reflex activity, we suggest that additional Ia and Ib discharges at long muscle lengths may enhance activation of the classical tri-synaptic pathway (Lalonde & Bui, 2021; Metz *et al*., 2023*b*). This could lead to an increase in post-activation depression by PAD, potentially reducing the contribution of Ia afferents to α-motoneuron discharge. However, it is important to consider that other mechanisms, particularly recurrent inhibition, are also likely to contribute to the overall increase in inhibition levels.

Our findings reveal a greater recurrent inhibition with increased muscle length (from short to long muscle length), regardless of contraction type. Considering our mechanical data demonstrating greater muscle fascicle length at long muscle length, coupled with the negative linear relationship between α-motoneuron discharge rate and the amount of recurrent inhibition (McCrea *et al*., 1980), it seems that the additional recurrent inhibition observed at longer muscle length may contribute to the reduction in α-motoneuron discharge rate. The hypothesis is supported by the work of Pasquet et al. (2006), which demonstrated that the discharge frequency of α-motoneurons is significantly higher when the muscle is shortened compared with when it is lengthened. However, these results were obtained on the tibialis anterior muscle, which may behave differently from the SOL muscle, particularly depending on the mode of contraction (Škarabot *et al*., 2019). Therefore, these data should be interpreted with caution. Given that the levels of descending neural drive, indicated by EMG activity, are comparable across the different muscle lengths, it seems unlikely that an enhancement due to descending neural drive is a contributing factor. Albeit speculative, it seems that the distinct behaviour of the motor unit at longer muscle lengths may be primarily attributed to the regulation of Renshaw cell activity rather than to a direct modulation of α-motoneuron activity by the descending neural drive. These observations are consistent with the findings of Hultborn *et al*. (1979), who demonstrated that descending neural drive provides a constant impulse directly to the α-motoneurons, whereas Renshaw cells (mediated by supraspinal and segmental influence) control the α-motoneuron frequency discharge. It is possible that Renshaw cells have a basal activity threshold specific to each muscle length, imposed by higher centres, which could be adjusted by the amount of descending neural drive and/or segmental afferent discharges (i.e. recurrent collaterals, afferents II and III). Furthermore, although the contribution of Ib inhibition cannot be entirely excluded (due to supramaximal stimulations), it is likely attenuated by the depolarisation of presynaptic inhibition at the Ib afferent terminals, which reduces Ib inhibitory interneurons activity during active movement (Lafleur *et al*., 1992; Özyurt *et al*., 2019). Consequently, it is unlikely that this mechanism underlies the modulations observed in our study.

## Conclusion

This study provides novel insights into how muscle length influences spinal inhibitory mechanisms during eccentric, isometric and concentric contractions. Our new findings reveal that increased muscle length reduces the effectiveness of Ia afferents to discharge α-motoneurons, primarily due to greater post-activation depression by PAD and/or greater recurrent inhibition activity. In addition, muscle length effect on post-activation depression by PAD was enhanced by eccentric contraction. However, although recurrent inhibition is more pronounced during eccentric contractions at intermediate and short muscle lengths relative isometric and concentric contractions, this is not true at long muscle lengths. At this muscle length the level of recurrent activity is similar between all contraction types. Despite the greater post-activation depression by PAD during eccentric contraction at long muscle length, we have shown that the effectiveness of activated Ia afferents to discharge α-motoneurons appears to vary similarly at recurrent inhibition activity. This work has revealed a previously unidentified function of recurrent inhibition that could potentially serve to modulate the frequency of α-motoneuron discharge in response to variations in muscle length.

## Data availability

Data will be available online at the time of publication.

## REFERENCES

1. Aagaard P, Simonsen EB, Andersen JL, Magnusson SP, Halkjaer-Kristensen J & Dyhre-Poulsen P (2000). Neural inhibition during maximal eccentric and concentric quadriceps contraction: effects of resistance training. J Appl Physiol (1985) 89, 2249–2257.

2. Aymard C, Katz R, Lafitte C, Lo E, Pénicaud A, Pradat-Diehl P & Raoul S (2000). Presynaptic inhibition and homosynaptic depression: a comparison between lower and upper limbs in normal human subjects and patients with hemiplegia. Brain 123 (Pt 8), 1688–1702.

3. Babault N, Pousson M, Michaut A & Van Hoecke J (2003). Effect of quadriceps femoris muscle length on neural activation during isometric and concentric contractions. Journal of Applied Physiology 94, 983–990.

4. Barrué-Belou S, Marque P & Duclay J (2018). Recurrent inhibition is higher in eccentric compared to isometric and concentric maximal voluntary contractions. Acta Physiol (Oxf) 223, e13064.

5. Barrué-Belou S, Marque P & Duclay J (2019). Supraspinal Control of Recurrent Inhibition during Anisometric Contractions. Med Sci Sports Exerc 51, 2357–2365.

6. Baudry S & Duchateau J (2012). Age-related influence of vision and proprioception on Ia presynaptic inhibition in soleus muscle during upright stance. J Physiol 590, 5541–5554.

7. Baudry S & Enoka RM (2009). Influence of load type on presynaptic modulation of Ia afferent input onto two synergist muscles. Exp Brain Res 199, 83–88.

8. Baudry S, Penzer F & Duchateau J (2014). Input-output characteristics of soleus homonymous Ia afferents and corticospinal pathways during upright standing differ between young and elderly adults. Acta Physiol (Oxf) 210, 667–677.

9. Bolsterlee B, Finni T, D’Souza A, Eguchi J, Clarke EC & Herbert RD (2018). Three-dimensional architecture of the whole human soleus muscle in vivo. PeerJ 6, e4610.

10. Burke D, Hagbarth KE & Löfstedt L (1978). Muscle spindle activity in man during shortening and lengthening contractions. J Physiol 277, 131–142.

11. Cattagni T, Merlet AN, Cornu C & Jubeau M (2018). H-reflex and M-wave recordings: effect of pressure application to the stimulation electrode on the assessment of evoked potentials and subject’s discomfort. Clin Physiol Funct Imaging 38, 416–424.

12. Colard J, Jubeau M, Crouzier M, Duclay J & Cattagni T (2024). Effect of muscle length on the modulation of H-reflex and inhibitory mechanisms of Ia afferent discharges during passive muscle lengthening. J Neurophysiol 132, 890–905.

13. Colard J, Jubeau M, Duclay J & Cattagni T (2023). Regulation of primary afferent depolarization and homosynaptic post-activation depression during passive and active lengthening, shortening and isometric conditions. Eur J Appl Physiol 123, 1257–1269.

14. Crone C & Nielsen J (1989). Methodological implications of the post activation depression of the soleus H-reflex in man. Exp Brain Res 78, 28–32.

15. Dimitriou M (2022). Human muscle spindles are wired to function as controllable signal-processing devices. Elife 11, e78091.

16. Doguet V, Nosaka K, Guével A, Thickbroom G, Ishimura K & Jubeau M (2017*a*). Muscle length effect on corticospinal excitability during maximal concentric, isometric and eccentric contractions of the knee extensors. Exp Physiol 102, 1513–1523.

17. Doguet V, Rivière V, Guével A, Guilhem G, Chauvet L & Jubeau M (2017*b*). Specific joint angle dependency of voluntary activation during eccentric knee extensions. Muscle Nerve 56, 750–758.

18. Duchateau J & Enoka RM (2016). Neural control of lengthening contractions. J Exp Biol 219, 197–204.

19. Duclay J & Martin A (2005). Evoked H-reflex and V-wave responses during maximal isometric, concentric, and eccentric muscle contraction. J Neurophysiol 94, 3555–3562.

20. Duclay J, Pasquet B, Martin A & Duchateau J (2011). Specific modulation of corticospinal and spinal excitabilities during maximal voluntary isometric, shortening and lengthening contractions in synergist muscles. J Physiol 589, 2901–2916.

21. Duclay J, Pasquet B, Martin A & Duchateau J (2014). Specific modulation of spinal and cortical excitabilities during lengthening and shortening submaximal and maximal contractions in plantar flexor muscles. J Appl Physiol (1985) 117, 1440–1450.

22. Funase K, Higashi T, Yoshimura T, Imanaka K & Nishihira Y (1996). Evident difference in the excitability of the motoneuron pool between normal subjects and patients with spasticity assessed by a new method using H-reflex and M-response. Neurosci Lett 203, 127–130.

23. Funase K, Imanaka K & Nishihira Y (1994). Excitability of the soleus motoneuron pool revealed by the developmental slope of the H-reflex as reflex gain. Electromyogr Clin Neurophysiol 34, 477–489.

24. Gerilovsky L, Tsvetinov P & Trenkova G (1989). Peripheral effects on the amplitude of monopolar and bipolar H-reflex potentials from the soleus muscle. Exp Brain Res 76, 173– 181.

25. Grosprêtre S, Papaxanthis C & Martin A (2014). Modulation of spinal excitability by a sub-threshold stimulation of M1 area during muscle lengthening. Neuroscience 263, 60–71.

26. Gruber M, Linnamo V, Strojnik V, Rantalainen T & Avela J (2009). Excitability at the Motoneuron Pool and Motor Cortex Is Specifically Modulated in Lengthening Compared to Isometric Contractions. Journal of Neurophysiology 101, 2030–2040.

27. Guilhem G, Doguet V, Hauraix H, Lacourpaille L, Jubeau M, Nordez A & Dorel S (2016). Muscle force loss and soreness subsequent to maximal eccentric contractions depend on the amount of fascicle strain in vivo. Acta Physiol (Oxf) 217, 152–163.

28. Haase J & van der meulen J (1961). Effects of supraspinal stimulation on Renshaw cells belonging to extensor motoneurones. J Neurophysiol 24, 510–520.

29. Hagood S, Solomonow M, Baratta R, Zhou BH & D’Ambrosia R (1990). The effect of joint velocity on the contribution of the antagonist musculature to knee stiffness and laxity. Am J Sports Med 18, 182–187.

30. Hari K, Lucas-Osma AM, Metz K, Lin S, Pardell N, Roszko DA, Black S, Minarik A, Singla R, Stephens MJ, Pearce RA, Fouad K, Jones KE, Gorassini MA, Fenrich KK, Li Y & Bennett DJ (2022). GABA facilitates spike propagation through branch points of sensory axons in the spinal cord. Nat Neurosci 25, 1288–1299.

31. Hopkins JT, Ingersoll CD, Cordova ML & Edwards JE (2000). Intrasession and intersession reliability of the soleus H-reflex in supine and standing positions. Electromyogr Clin Neurophysiol 40, 89–94.

32. Howatson G, Taylor MB, Rider P, Motawar BR, McNally MP, Solnik S, DeVita P & Hortobágyi T (2011). Ipsilateral motor cortical responses to TMS during lengthening and shortening of the contralateral wrist flexors. Eur J Neurosci 33, 978–990.

33. Hultborn H, Lindström S & Wigström H (1979). On the function of recurrent inhibition in the spinal cord. Exp Brain Res 37, 399–403.

34. Hultborn H, Meunier S, Pierrot-Deseilligny E & Shindo M (1987). Changes in presynaptic inhibition of Ia fibres at the onset of voluntary contraction in man. J Physiol 389, 757–772.

35. Hwang IS (2002). Assessment of soleus motoneuronal excitability using the joint angle dependent H reflex in humans. J Electromyogr Kinesiol 12, 361–366.

36. Jankowska E (1992). Interneuronal relay in spinal pathways from proprioceptors. Prog Neurobiol 38, 335–378.

37. Jankowska E & Edgley SA (2010). Functional subdivision of feline spinal interneurons in reflex pathways from group Ib and II muscle afferents; an update. Eur J Neurosci 32, 881– 893.

38. Johannsson J, Duchateau J & Baudry S (2015). Presynaptic inhibition of soleus Ia afferents does not vary with center of pressure displacements during upright standing. Neuroscience 298, 63–73.

39. Katz & Pierrot-Deseilligny E (1999). Recurrent inhibition in humans. Prog Neurobiol 57, 325–355.

40. Kidgell DJ, Bonanno DR, Frazer AK, Howatson G & Pearce AJ (2017). Corticospinal responses following strength training: a systematic review and meta-analysis. Eur J Neurosci 46, 2648–2661.

41. Klimstra M & Zehr EP (2008). A sigmoid function is the best fit for the ascending limb of the Hoffmann reflex recruitment curve. Exp Brain Res 186, 93–105.

42. Komi PV, Linnamo V, Silventoinen P & Sillanpää M (2000). Force and EMG power spectrum during eccentric and concentric actions. Med Sci Sports Exerc 32, 1757–1762.

43. Lafleur J, Zytnicki D, Horcholle-Bossavit G & Jami L (1992). Depolarization of Ib afferent axons in the cat spinal cord during homonymous muscle contraction. The Journal of Physiology 445, 345–354.

44. Lalonde NR & Bui TV (2021). Do spinal circuits still require gating of sensory information by presynaptic inhibition after spinal cord injury? Current Opinion in Physiology 19, 113– 118.

45. Lamy J-C, Wargon I, Mazevet D, Ghanim Z, Pradat-Diehl P & Katz R (2009). Impaired efficacy of spinal presynaptic mechanisms in spastic stroke patients. Brain 132, 734–748.

46. Magalhães FH, Elias LA, da Silva CR, de Lima FF, de Toledo DR & Kohn AF (2015). D1 and D2 Inhibitions of the Soleus H-Reflex Are Differentially Modulated during Plantarflexion Force and Position Tasks. PLoS One 10, e0143862.

47. Marchand-Pauvert V, Nicolas G, Marque P, Iglesias C & Pierrot-Deseilligny E (2005). Increase in group II excitation from ankle muscles to thigh motoneurones during human standing. The Journal of Physiology 566, 257–271.

48. Matthews PBC (2011). Muscle Spindles: Their Messages and Their Fusimotor Supply. In Comprehensive Physiology, pp. 189–228. John Wiley & Sons, Ltd. Available at: https://onlinelibrary.wiley.com/doi/abs/10.1002/cphy.cp010206 [Accessed November 16, 2022].

49. McCrea DA, Pratt CA & Jordan LM (1980). Renshaw cell activity and recurrent effects on motoneurons during fictive locomotion. J Neurophysiol 44, 475–488.

50. Metz K, Matos IC, Hari K, Bseis O, Afsharipour B, Lin S, Singla R, Fenrich KK, Li Y, Bennett DJ & Gorassini MA (2023*a*). Post-activation depression from primary afferent depolarization (PAD) produces extensor H-reflex suppression following flexor afferent conditioning. J Physiol 601, 1925–1956.

51. Metz K, Matos IC, Li Y, Afsharipour B, Thompson CK, Negro F, Quinlan KA, Bennett DJ & Gorassini MA (2023*b*). Facilitation of sensory transmission to motoneurons during cortical or sensory-evoked primary afferent depolarization (PAD) in humans. J Physiol 601, 1897–1924.

52. Meunier S, Pierrot-Deseilligny E & Simonetta-Moreau M (1994). Pattern of heteronymous recurrent inhibition in the human lower limb. Exp Brain Res 102, 149–159.

53. Mizuno Y, Tanaka R & Yanagisawa N (1971). Reciprocal group I inhibition on triceps surae motoneurons in man. J Neurophysiol 34, 1010–1017.

54. Nordez A, Cornu C & McNair P (2006). Acute effects of static stretching on passive stiffness of the hamstring muscles calculated using different mathematical models. Clin Biomech (Bristol, Avon) 21, 755–760.

55. Özyurt MG, Piotrkiewicz M, Topkara B, Weisskircher H-W & Türker KS (2019). Motor units as tools to evaluate profile of human Renshaw inhibition. J Physiol 597, 2185–2199.

56. Papitsa A, Paizis C, Papaiordanidou M & Martin A (2022). Specific modulation of presynaptic and recurrent inhibition of the soleus muscle during lengthening and shortening submaximal and maximal contractions. J Appl Physiol (1985) 133, 1327–1340.

57. Pasquet B, Carpentier A & Duchateau J (2006). Specific modulation of motor unit discharge for a similar change in fascicle length during shortening and lengthening contractions in humans. J Physiol 577, 753–765.

58. Pasquet B, Carpentier A, Duchateau J & Hainaut K (2000). Muscle fatigue during concentric and eccentric contractions. Muscle Nerve 23, 1727–1735.

59. Penzer F, Duchateau J & Baudry S (2015). Effects of short-term training combining strength and balance exercises on maximal strength and upright standing steadiness in elderly adults. Exp Gerontol 61, 38–46.

60. Petersen NT, Butler JE, Carpenter MG & Cresswell AG (2007). Ia-afferent input to motoneurons during shortening and lengthening muscle contractions in humans. Journal of Applied Physiology 102, 144–148.

61. Pierrot-Deseilligny E & Burke D (2005). The Circuitry of the Human Spinal Cord: Its Role in Motor Control and Movement Disorders. Cambridge University Press.

62. Pierrot-Deseilligny E & Bussel B (1975). Evidence for recurrent inhibition by motoneurons in human subjects. Brain Res 88, 105–108.

63. Rudomin P & Schmidt RF (1999). Presynaptic inhibition in the vertebrate spinal cord revisited. Exp Brain Res 129, 1–37.

64. Schieppati M (1987). The Hoffmann reflex: a means of assessing spinal reflex excitability and its descending control in man. Prog Neurobiol 28, 345–376.

65. Simonetta-Moreau M, Marque P, Marchand-Pauvert V & Pierrot-Deseilligny E (1999). The pattern of excitation of human lower limb motoneurones by probable group II muscle afferents. The Journal of Physiology 517, 287–300.

66. Škarabot J, Ansdell P, Brownstein CG, Hicks KM, Howatson G, Goodall S & Durbaba R (2019). Corticospinal excitability of tibialis anterior and soleus differs during passive ankle movement. Exp Brain Res 237, 2239–2254.

67. Souron R, Baudry S, Millet GY & Lapole T (2019). Vibration-induced depression in spinal loop excitability revisited. J Physiol 597, 5179–5193.

68. Theodosiadou A, Henry M, Duchateau J & Baudry S (2023). Revisiting the use of Hoffmann reflex in motor control research on humans. Eur J Appl Physiol 123, 695–710.

69. Westing SH, Cresswell AG & Thorstensson A (1991). Muscle activation during maximal voluntary eccentric and concentric knee extension. Eur J Appl Physiol Occup Physiol 62, 104–108.

70. Windhorst U (1996). On the role of recurrent inhibitory feedback in motor control. Prog Neurobiol 49, 517–587.

